# Geographic potential of the world’s largest hornet, *Vespa mandarinia* Smith (Hymenoptera: Vespidae), worldwide and particularly in North America

**DOI:** 10.1101/2020.08.11.246991

**Authors:** Claudia Nuñez-Penichet, Luis Osorio-Olvera, Victor H. Gonzalez, Marlon E. Cobos, Laura Jiménez, Devon A. DeRaad, Abdelghafar Alkishe, Rusby G. Contreras-Díaz, Angela Nava-Bolaños, Kaera Utsumi, Uzma Ashraf, Adeola Adeboje, A. Townsend Peterson, Jorge Soberón

**Author notes:** Corresponding Author: Jorge Soberón, Department of Ecology & Evolutionary Biology and Biodiversity Institute, University of Kansas, 1345 Jayhawk Blvd., Lawrence, Kansas 66045, USA.

## Abstract

The Asian giant hornet (AGH, *Vespa mandarinia*) is the world’s largest hornet, occurring naturally in the Indomalayan region, where it is a voracious predator of pollinating insects including honey bees. In September 2019, a nest of Asian giant hornets was detected outside of Vancouver, British Columbia and in May 2020 an individual was detected nearby in Washington state, indicating that the AGH successfully overwintered in North America. Because hornets tend to spread rapidly and become pests, reliable estimates of the potential invasive range of *V. mandarinia* in North America are needed to assess likely human and economic impacts, and to guide future eradication attempts. Here, we assess climatic suitability for AGH in North America, and suggest that, without control, this species could establish populations across the Pacific Northwest and much of eastern North America. Predicted suitable areas for AGH in North America overlap broadly with areas where honey production is highest, as well as with species-rich areas for native bumble bees and stingless bees of the genus *Melipona* in Mexico, highlighting the economic and environmental necessity of controlling this nascent invasion.

## Introduction

Invasive species represent major threats to biodiversity, as they can alter ecosystem processes and functions (Pyšek & Richardson, 2010; Vilà et al., 2011), and often contribute to the decline of imperiled species (e.g., Wilcove et al., 1998; Dueñas et al., 2018). The economic damage to agriculture, forestry, and public health, resulting from invasive species totals nearly $120 billion annually in the United States alone (Pimentel, Zuniga & Morrison, 2005).

Even in the midst of the global uncertainty and socio-economic distress resulting from the COVID-19 pandemic, the recent detection of the Asian Giant Hornet (AGH, *Vespa mandarinia* Smith, Hymenoptera: Vespidae), in North America (Bérubé, 2020), received significant public attention. This social insect is the world’s largest hornet (2.5–4.5 cm body length), and occurs naturally across Asia, including in India, Nepal, Sri Lanka, Vietnam, Taiwan, and Japan, at elevations ranging between 850 and 1900 (Matsuura & Sakagami, 1973; Archer, 2008; Smith-Pardo, Carpenter & Kimsey, 2020). As in other temperate-zone social species, annual colonies of the AGH, which may contain up to 500 workers, die at the onset of winter and mated queens overwinter in underground cavities. After emerging in the spring, each queen starts a new colony in a pre-existing cavity, typically in tree roots or an abandoned rodent nest (Archer, 2008). Like other species of *Vespa*, AGH is a voracious predator of insects, particularly honey bees and other social wasps. Attacks on honey bee hives occur late in the development of the hornet colony and prior to the emergence of reproductive individuals (males and new queens), the timing of which depends on location (e.g., Matsuura & Sakagami, 1973; Matsuura, 1988; Archer, 2008).

In its native range, AGH attacks several species of bees, some of which have developed sophisticated defense mechanisms against attacks (Ono et al., 1995; Kastberger, Schmelzer & Kranner, 2008; Fujiwara, Sasaki & Washitani, 2016). The best documented, colony-level defense mechanism is in the Asiatic honey bee, *Apis cerana* Fabricius, which can detect site-marking pheromones released by AGH scouts, and responds by engulfing a single hornet in a ball consisting of up to 500 bees. The heat generated by the vibration of the bees’ flight muscles, and the resulting high levels of CO2 from respiration effectively kill the hornet (Ono et al., 1995; Sugahara, Nishimura & Sakamoto, 2012). In contrast, European honey bees (*A. mellifera* L.) cannot detect and respond to AGH marking pheromones and colonies are more or less defenseless against AGH attacks (McClenaghan et al., 2019). As few as a dozen AGH can destroy a European honey bee colony of up to 30,000 individuals (Matsuura & Sakagami, 1973).

In addition to the threat to the beekeeping industry, the introduction of AGH in North America is also concerning for public health. Their powerful stings can induce severe allergic reactions or even death in hypersensitive individuals (Schmidt et al., 1986; Yanagawa et al., 2007). Annually, 30-40 people die from AGH stings in Japan, most as a result of anaphylaxis or sudden cardiac arrest (Matsuura & Sakagami, 1973); similar deadly cases have been reported from China (Li et al., 2015).

Although invasive species are typically limited by dispersal ability and suitability of novel environments, vespid hornets are well known for their invasive success and excellent dispersal capacity (Beggs et al., 2011; Monceau, Bonnard & Thiéry, 2014). As such, the introduction of AGH in the Pacific Northwest poses a potentially serious ecological and socio-economic risk in North America. Here, we use ecological niche modeling (ENM) to detect areas of suitable environments for this species worldwide, with particular emphasis on North America. We also use a dispersal simulation approach to detect potential invasion paths of this species within North America. A similar methodology for projecting AGH invasion potential has been implemented by Zhu et al. (2020); we build upon this framework by introducing several modifications to the modelling approach, and investigating further the potential ecological and economic impacts of an AGH invasion in North America.

## Methods

### Occurrence and environmental data

We downloaded occurrence data for *V. mandarinia* from the Global Biodiversity Information Facility database (GBIF; https://www.gbif.org/). We kept records from the species’ native range (Fig. 1) separate from non-native occurrences facilitated by human introduction. We cleaned occurrences from the native distribution following Cobos et al. (2018) by removing duplicates and records with doubtful or missing coordinates. To avoid model overfitting derived from spatial autocorrelation and overdominance of specific regions due to sampling bias, we thinned these records spatially in two ways: by geographic distance and by density of records per country (Fig. 2). In the first case (distance-based thinning), we excluded occurrences that were <50 km away from another locality (Anderson, 2012). In the second thinning approach (country-density thinning), we randomly reduced numbers of occurrences in countries with the densest sampling, namely Japan, Taiwan, and South Korea (from 30, 6, and 5, to 6, 2, and 2 occurrences, respectively), to match an approximate reference density of India, Nepal, and China. We used the package ellipsenm (Cobos et al., 2020; available at https://github.com/marlonecobos/ellipsenm) in R 3.6.2 (R Core Team, 2019) to clean and thin the data. We then treated both data sets independently in all subsequent analysis steps.

**Figure 1.**
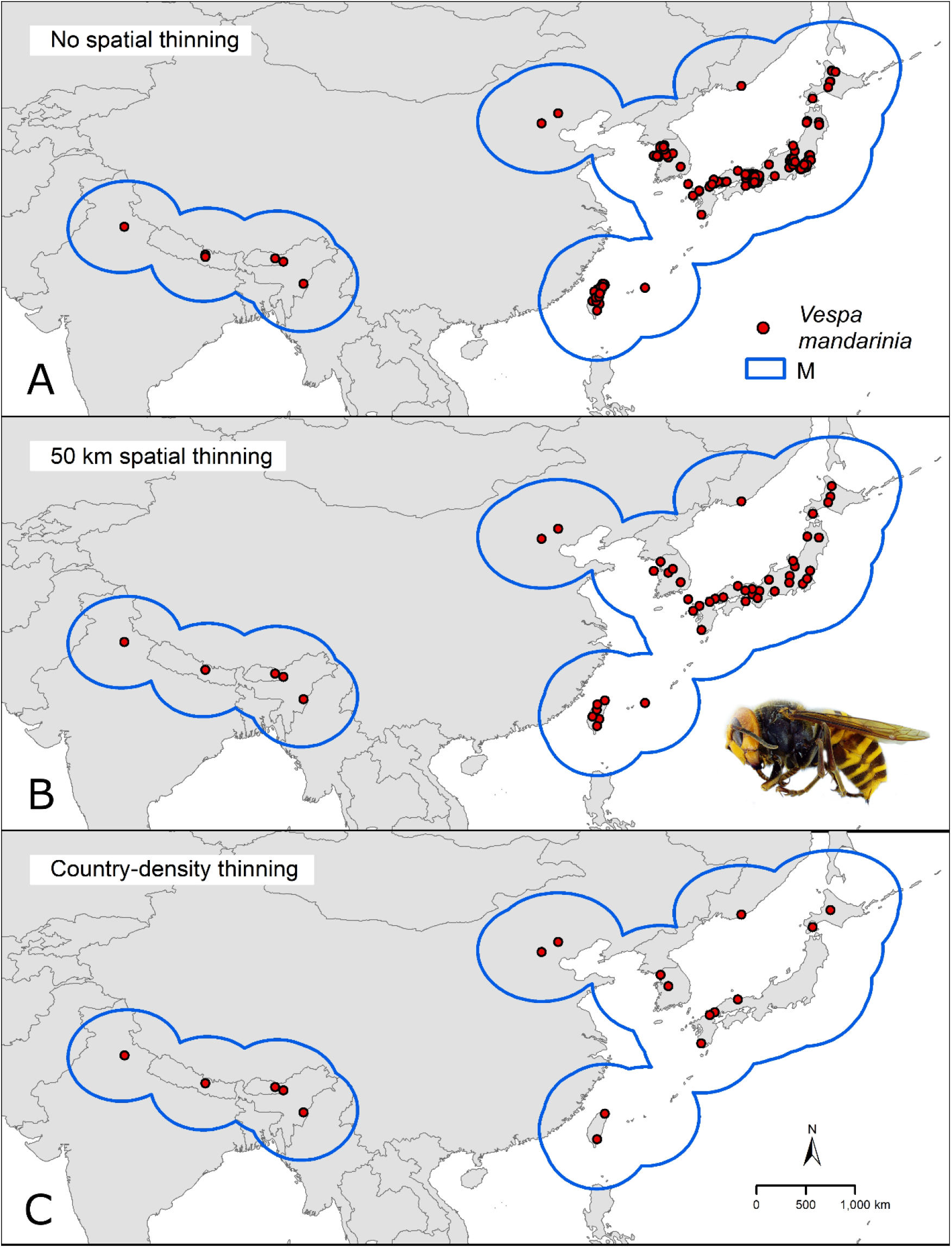
Hypothesis of accessible areas (**M**) and occurrence records of *Vespa mandarinia* across its native distribution. The three panels represent the occurrences left after cleaning (A) and after applying the two thinning approaches (B and C).

**Figure 2.**
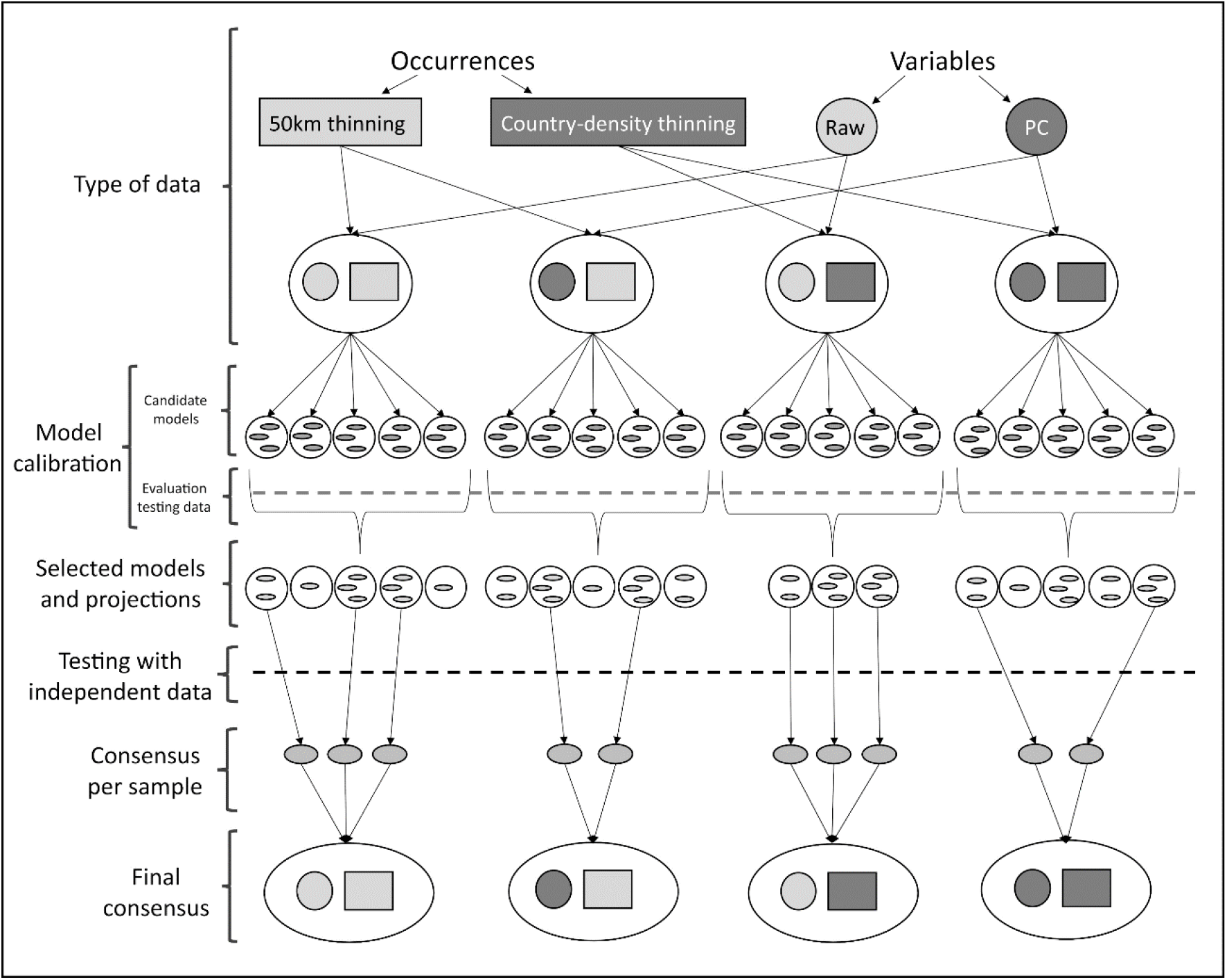
Schematic representation of methods used to obtain ecological niche models for *Vespa mandarinia*, considering the uncertainty coming from the distinct treatments applied to the data and the variability resulting from different procedures and methodological decisions made during model calibration.

For environmental predictors, we used bioclimatic variables at 10’ resolution (~18 km at the Equator) from the MERRAclim database (Vega, Pertierra & Olalla-Tárraga, 2018). We excluded four variables because they are known to contain spatial artifacts as a result of combining temperature and humidity information (Escobar et al., 2014): mean temperature of most humid quarter, mean temperature of least humid quarter, specific humidity mean of warmest quarter, and specific humidity mean of coldest quarter. The 15 variables remaining were masked to an area for model calibration (**M**, see Ecological niche modeling).

These 15 variables were submitted to a principal component analysis (PCA) to reduce dimensionality and multicollinearity. Raw variables and principal components (PCs) were considered separately in all subsequent analyses. To select a set of raw variables, we reduced them to a subset with Pearson’s correlation coefficients (*r*) ≤ 0.85, choosing the most biologically relevant or interpretable variables based on our knowledge of AGH natural history (Simões et al., 2020). The PCA was calibrated using environmental variation across the **M** area, and transferred to the whole world. All analyses were done in R; specifically, raster processing was done using the packages raster (Hijmans et al., 2020), rgeos (Bivand et al., 2020b), and rgdal (Bivand et al., 2020a); PCA was done using the ntbox package (Osorio-Olvera et al., 2020; available in https://github.com/luismurao/ntbox).

### Ecological niche modeling

To identify a calibration area (ostensibly equivalent to **M**; Owens et al., 2013) for our models, we considered a region contained within a buffer of 500 km around the known occurrence records after the 50 km thinning process (Fig. 1). This distance was selected considering the species’ dispersal ability (Matsuura & Sakagami, 1973; APHIS, 2020). We used all pixels in **M** (15,411) as the background across which to calibrate the models.

Given uncertainty deriving from specific treatments of occurrence records and environmental predictors in ecological niche modeling (Alkishe et al., 2020), we calibrated models via four distinct schemes: (1) using raw variables and distance-based thinned occurrences, (2) using PCs and distance-based thinned occurrences, (3) using raw environmental variables and country-density thinned occurrences, and (4) using PCs and country-density thinned occurrences (Fig. 2). For each scheme, we calibrated models five times, each time randomly selecting 50% of the occurrences for calibrating models, and using the remaining records for testing (Cobos et al., 2019a).

Each process of model calibration consisted of creating and evaluating candidate models using Maxent (Phillips, Anderson & Schapire, 2006; Phillips et al., 2017) based on distinct parameter settings: 10 regularization multiplier values (0.10, 0.25, 0.50, 0.75, 1, 2, 3, 4, 5, 6), eight feature classes (lq, lp, lqp, qp, q, lqpt, lqpth, lqph, where l is linear, q is quadratic, p is product, t is threshold, and h is hinge), and all combinations of more than two predictor variables (Cobos et al., 2019b; Table S1-S2). We tested 4560 models using raw variables and 880 using PCs (see Data preprocessing and model calibration), in tandem with the two methods of reducing occurrence data described above. We assessed model performance using partial ROC (for statistical significance; Peterson, Papeş & Soberón, 2008), omission rates (*E* = 5%, for predictive ability; Anderson, Lew & Peterson, 2003), and Akaike Information Criterion corrected for small sample sizes (AICc, for model complexity; Warren & Seifert, 2011). We selected models with delta AICc ≤2 (Cobos et al., 2019a) from those that were statistically significant and had omission rates below 5%.

After model calibration, we created models with the selected parameter values, using all occurrences after the corresponding thinning process, with 10 bootstrap replicates, cloglog output, and model transfers using three types of extrapolation (free extrapolation, extrapolation and clamping, no extrapolation; Owens et al., 2013). Not all calibration processes identified models that met all three criteria of model selection; we did not consider those models in further analyses (Fig. 2; Table 1). As a final evaluation step, we tested whether each replicate of the selected models was able to anticipate the known invasive records of the species in the Americas (British Columbia, Canada; Washington, USA). For each scheme, using only those model replicates that met the selection criteria and correctly predicted independent occurrences, we created two types of consensus: (1) a median of the medians obtained for each parameterization, and (2) the sum of all suitable areas derived from binarizing each replicate using a modified least presence (5% omission) threshold (Fig. 2).

**Table 1.**
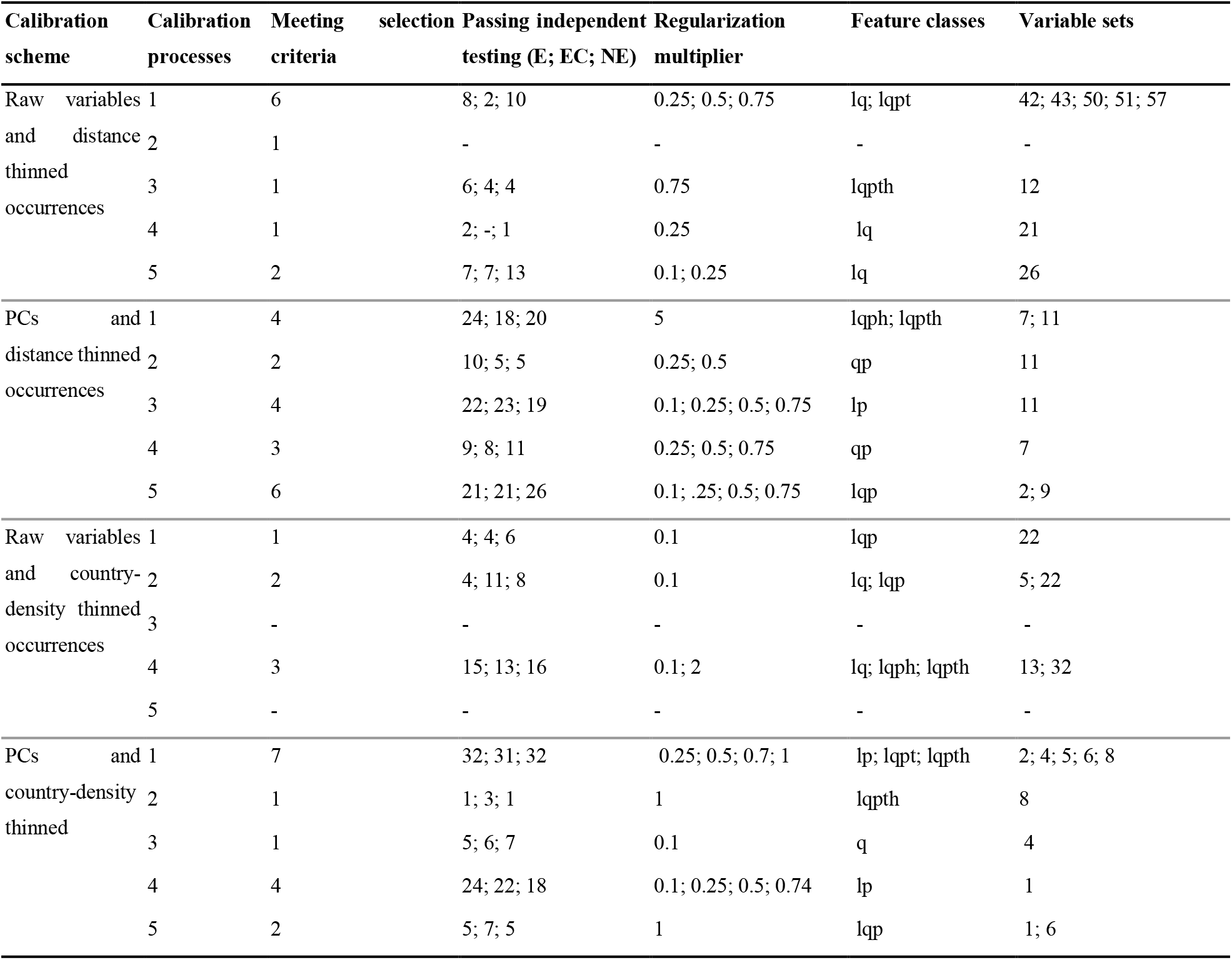
Summary of results of ecological niche modeling for *Vespa mandarinia*, including model calibration, model evaluation, and features for models selected after independent testing. The variables included in the sets mentioned on this table can be found in Table 1S-2S. E: free extrapolation; EC: extrapolation and clamping; NE: no extrapolation.

As we transferred models to the entire world, we used the mobility-oriented parity metric (MOP; Owens et al., 2013) to detect areas where strict or combinational extrapolation risks could be expected, given the presence of non-analogous conditions with respect to the environments manifested across the calibration area. We used areas where extrapolation risks were detected using MOP to trim our binary results (suitable areas) to avoid potentially problematic interpretations based on extrapolative situations. Model calibration, production of selected models with replicates, and MOP analyses were done in R using the package kuenm (Cobos et al., 2019a); raster processing and independent testing of models were done using the package raster and other base functions in R.

### Dispersal simulations

We used the binary outputs from the final consensus models (suitable and unsuitable areas, without areas of strict extrapolation) to simulate invasion dynamics of the AGH. All simulations were started from the Pacific Northwest, from sites already known to be occupied by the AGH. The simulations were performed using the cellular automaton dynamic model included in the bam R package (Osorio-Olvera & Soberón, 2020; available at https://github.com/luismurao/bam). Under this discrete model, given an occupied area at time *t*, two layers of information are needed to obtain the occupied area at time *t +1:* (i) the binary layer of suitability for the species, and (ii) a connectivity matrix determined by the species’ ability to reach neighboring cells in one time unit (known as “Moore’s neighborhood”; Gray et al., 2003, that defines patches that are connected by dispersal). At each step, each of the suitable cells can be either occupied or not by the species. If a cell is occupied, adjacent cells can be visited by the species, and if suitable, they become occupied. This method is similar to the one implemented in the MigClim R package (Engler, Hordijk & Guisan, 2012), but uses a simpler dispersal kernel and parameterization.

For each of the schemes followed to obtain ecological niche models for *V. mandarinia*, we performed a set of simulations in which we explored different degrees of connectivity (1–8, 10, and 12 neighbor cells) and different suitability thresholds (10 equidistant levels from 3–10% of the presence points) to create the binary maps. All simulations were done with 200 steps. In the end, we visualized the simulation results by summing the occupied distribution layers obtained from each set of simulations. A value of 100 in these final layers means that the species reached that cell in 100% of the simulations, whereas a value of 0 means that the species never reached that cell. Further details regarding the simulation processes can be found in the Supplementary Information.

### Honey production and native bee richness in North America

To explore potential ecological and economic impacts of the invasion of the AGH in North America, we explored annual, state-level production of honey as well as species richness of bumble bees (*Bombus* Latreille) and stingless bees (*Melipona* Illiger) in Mexico and the United States. We extracted data on 2016 honey production (in US dollars) for the United States from the U.S. Department of Agriculture (USDA; available at https://quickstats.nass.usda.gov/#4A0314DA-F3E5-3B06-ADD1-CA8032FBD937) and from the Instituto Nacional de Estadística, Geografía e Informática (INEGI) for Mexico (https://atlasapi2019.github.io/cap4.html). For native species richness, we obtained a list of species of bumble bees and stingless bees that occur in Mexico and the United States from Discover Life (https://www.discoverlife.org/) and downloaded their occurrence data from GBIF. We chose these bee taxa as likely targets of AGH because the species in these groups are of similar body size and behavior to the typical prey of these hornets: they are social insects that form annual or perennial colonies that can have a few hundreds to as many as 10,000 individuals (Cueva del Castillo, Sanabria-Urbán & Serrano-Meneses, 2015; Viana et al., 2015), and store honey and pollen inside their nests (Michener, 2000). To summarize species richness of these two genera, we created a presence absence matrix (PAM; Arita et al., 2008) for North America, based on geographic coordinates of occurrence data, with a pixel size of one degree. The PAM was created in R with the package biosurvey (Nuñez-Penichet et al., 2020; available at https://github.com/claununez/biosurvey).

To assure transparency and reproducibility of our work, we include an Overview, Data, Model, Assessment, and Prediction protocol (ODMAP; Zurell et al., 2020) in our supplementary materials. This metadata summary provides a detail key steps included in our analyses.

## Results

### Data preprocessing and model calibration

We retained 172 occurrence records for *V. mandarinia* after initial data cleaning, 49 records after the distance-based thinning approach, and 18 records after the country-density thinning approach (Fig. 1). As environmental predictors, we selected six raw variables based on correlation levels and natural history criteria: isothermality (BIO3), maximum temperature of warmest month (BIO5), minimum temperature of coldest month (BIO6), temperature annual range (BIO7), specific humidity of most humid month (BIO13), and specific humidity of least humid month (BIO14). From the PCA, we kept the first four PC axes, as they explained 97.9% of the cumulative variance (Figure S1).

The number of models that met the selection criteria was considerably smaller than the total number of models tested (Table 1). The calibration schemes including raw variables had fewer models selected than the those using PCs (11, 19, 6, 15 models selected for raw/distance-thinned, PC/distance-thinned, raw/country-density, and PC/country-density, respectively). The number of replicates of those selected models that predicted the *V. mandarinia* invaded areas in North America was also small and changed among types of extrapolation (Table 1).

### Ecological niche model predictions

In our models, areas predicted as suitable for the AGH varied among calibration schemes, in both scale and geographic pattern (Fig. 3, Figures S2-S4). The differences are conspicuous between the two types of thinning approaches, which resulted in models with different numbers of occurrence records. Models with country-density thinning (18 records) resulted in broad predicted suitable areas worldwide, with areas of higher values of suitability concentrated in tropical regions (Fig. 3, Figures S2-S4). In contrast, models created with the greater number of occurrences (49 records) from the geographic distance thinning predicted more patches of suitable areas across large extensions of Southeast Asia, Europe, West Africa, Central America, northern South America, and the Pacific Northwest and southeastern United States (Fig. 3, Figures S2-S4). In the calibration area, the areas detected with high levels of suitability were larger in the scheme with geographic distance thinned occurrences and the raw variables and smaller in the predictions obtained with the country-density thinned occurrences and the PCs as environmental predictors (Fig. 3). In all schemes, the two northernmost occurrence points of this species in China were accorded relatively low levels of suitability (Fig. 3). Predicted suitable areas for this hornet worldwide were also different among types of extrapolation considered in this study, especially as regards distribution size rather than location (Figures S2-S4).

**Figure 3.**
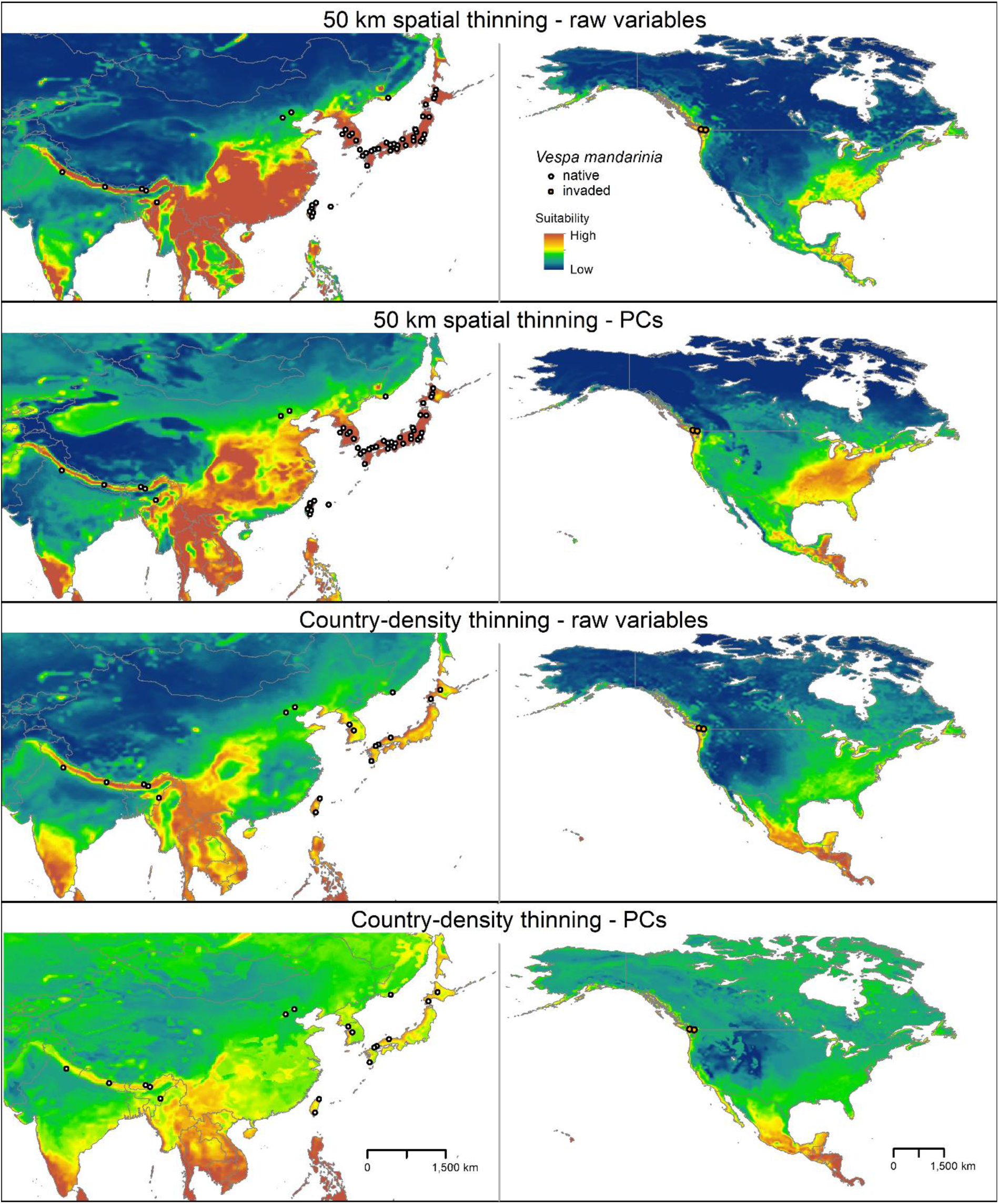
Median of potentially suitable areas for *Vespa mandarinia* predicted with free extrapolation for different calibration schemes in the calibration area (left panels) and in North America (right panels).

In North America, across multiple model calibration schemes, our various models agreed in predicting suitable areas for AGH in the Pacific region of southwestern Canada, the Pacific Northwest, the southeastern United States, and from central Mexico south to southernmost Panama (Fig. 4). Our model calibration schemes also agreed in identifying the Rocky Mountains and Great Plains as unsuitable for this species (Fig. 4).

**Figure 4.**
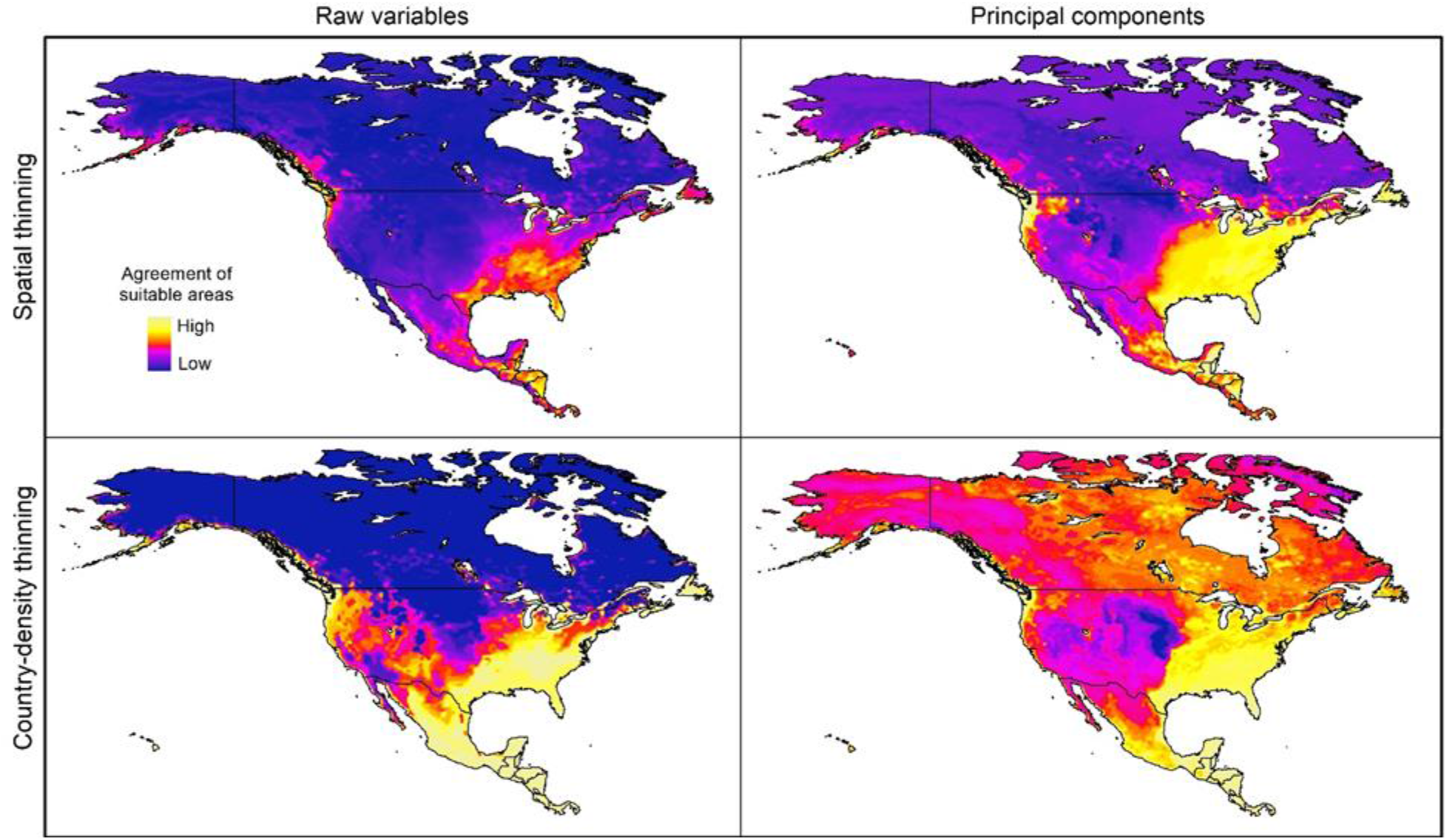
Sum of all suitable areas for *Vespa mandarinia* in North America derived from binarizing (using a 5% threshold) each replicate of selected models (model transfers done with extrapolation) that predicted the known invaded localities of this hornet.

Prevalences (proportion of suitable area) varied among the data thinning schemes. In the case of models created with raw variables, prevalence values of 0.171 and 0.164 were detected when using spatially rarefied and country-density rarefied records, respectively. Models created with PCs had prevalences of 0.248 and 0.239 for spatially rarefied and country-density rarefied records, respectively (Table S3).

### Extrapolation risks in model projections

The pattern of areas detected with risk of extrapolation was similar worldwide between thinning methods, but different between raw variables and PCs (Fig. 5, Figure S8). Most tropical areas predicted as suitable were identified as regions with high extrapolation risk (Figure S8). For raw variables, the areas with extrapolation risk in North America included most of Canada and Alaska, whereas for PCs areas with extrapolation risk included large portions of Mexico and, the central-southwestern United States, as well as the islands north of Hudson Bay in Canada (Fig. 5).

**Figure 5.**
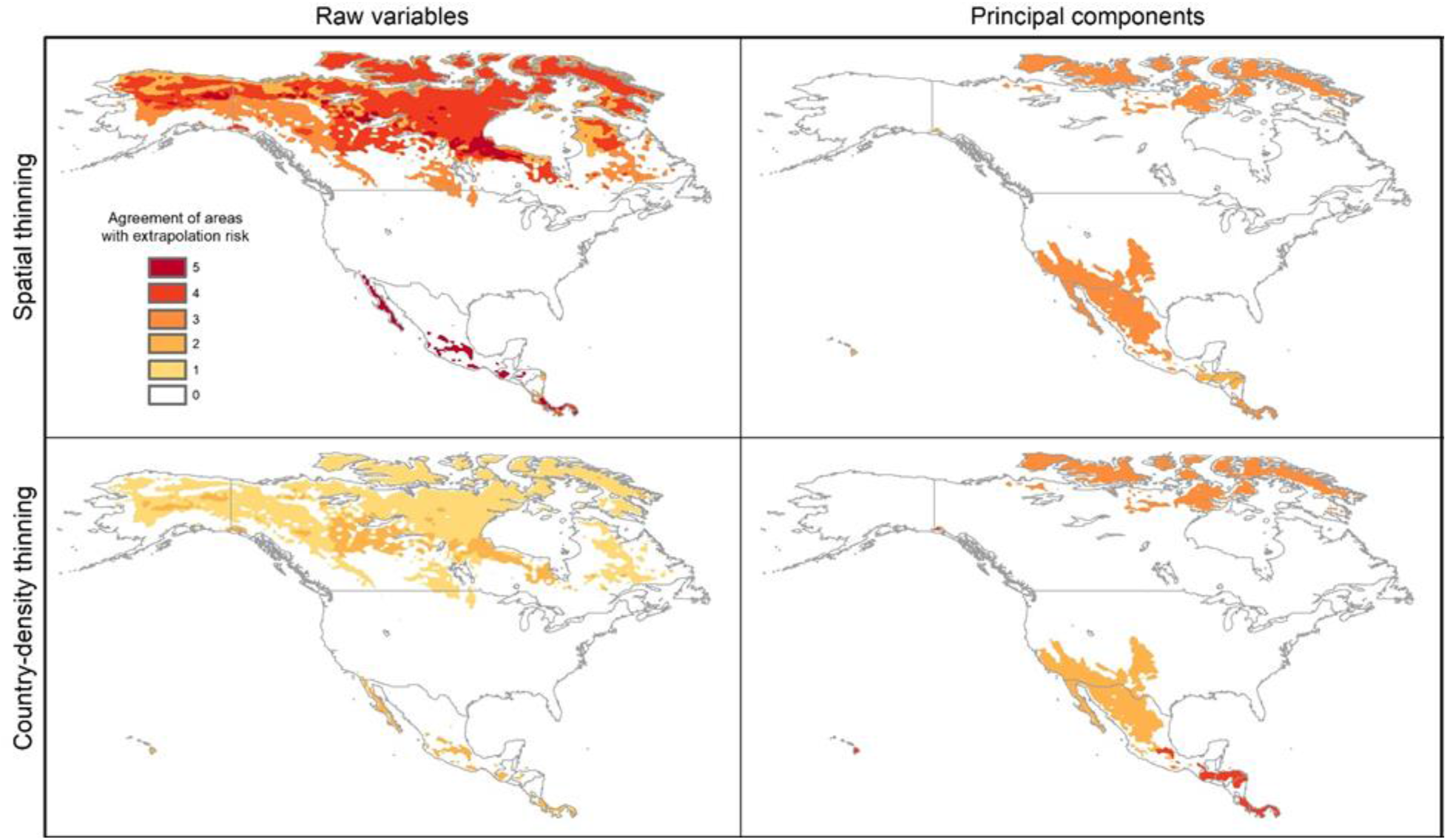
Agreement of areas with extrapolation risk for models of *Vespa mandarinia* in North America, separated by calibration schemes.

### Simulations of potential invasion

The simulations of potential sequences of colonization and dispersal of AGH in North America, starting from the known invaded localities, showed agreement among calibration schemes in predicting an invasion across the Pacific Northwest from southernmost Alaska to southernmost California in the United States (Fig. 6). In contrast, we found that the dispersal distance required to invade all the way to the East Coast of North America varied among calibration schemes. In the schemes using raw variables, the route of invasion to reach the East Coast goes from the Pacific Northwest down to California and Mexico, and then up the East Coast of North America. A dispersal distance of 10 cells (where each cell represents ~18 km) was enough to reach the East Coast (see left panels in Fig. 6). For the scheme using the 50 km spatially-thinned occurrences and PCs, the invasion follows a more direct route from the Pacific Northwest to the East Coast that goes through the United States, and the required dispersal distance to reach the East Coast was only 4 cells (top right panel in Fig. 6). Finally, in the case of country-density thinned occurrences and PCs, the invasion goes from the Pacific Northwest through Canada to the Atlantic, and then down the East Coast to the United States. A distance of 8 cells was needed to make this invasion route possible (bottom right panel in Fig. 6).

**Figure 6.**
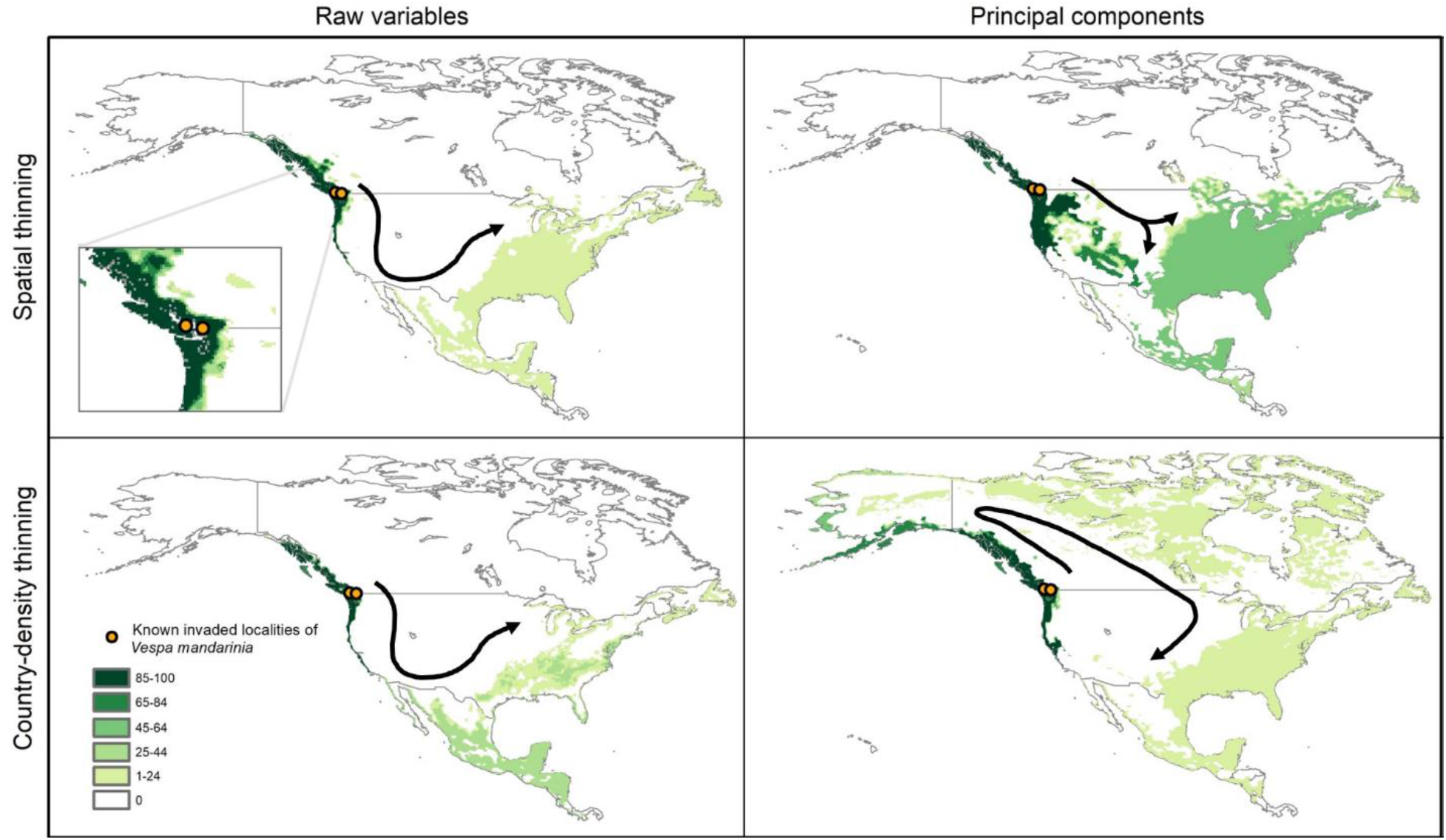
Results from simulations of the potential dynamics of invasion of *Vespa mandarinia* in North America. Dark shades of green show areas that the species reached in a high percentage of scenarios, while light shades of green represent areas reached only rarely by the species. Arrows represent the general path of potential invasion.

## Discussion

The patterns of suitability that we found in North America across multiple input data processing schemes (Fig. 6) are broadly concurrent with the results obtained by Zhu et al. (2020), who used an ensemble modeling approach. This concordance (both among our selected models, and between our models and the ensemble models), gives us confidence that the Pacific Northwest and southeastern United States represent suitable areas for AGH. In contrast with the results of Zhu et al. (2020), however, our dispersal simulations indicate a larger potential invasion area in the United States, with the AGH potentially crossing to eastern North America via a southern invasion route, through Mexico and Texas; a southeast-ward route crossing Idaho, Wyoming, and Colorado; or a northern route across Canada and the Great Lakes region (Fig. 6).

Quantifying the probability of the AGH following any one of the individual dispersal routes presented would require precise quantification of dispersal ability, and discerning the real-world validity of each of the four modeling outcomes. Instead of attempting to guess, we present several models that offer multiple plausible invasion scenarios. Across all scenarios presented, the AGH is expected to establish populations along the coastal Pacific Northwest via short-distance dispersal, and it is likely to invade the southeastern United States if it has even moderate dispersal potential (Fig. 6). It is important to note that these potential invasion routes consider only the natural dispersal ability of this hornet, and do not take into account the effect of potential accidental human-aided dispersal through the transport of soil and wood, where fertilized queen AGHs overwinter (Archer, 1995). Such unwitting human-aided dispersal is a serious concern, as it could potentiate a rapid invasion of this hornet to environmentally suitable, yet currently isolated places across North America. Our simulations with larger numbers for neighbor cells are perhaps a good illustration of what could be expected if dispersal events to very long distances occur.

Contrasts between our prediction of extensive invasion potential, and Zhu et al.’s (2020) more conservative predictions, arise from Zhu et al.’s (2020) use of MigClim (Engler, Hordijk & Guisan, 2012) to model dispersal of the AGH in western North America. MigClim is a cellular automaton platform that models the state of grid cells as occupied or unoccupied. Although we used the same modelling technique, our dispersal kernel is a much simpler “Moore Neighborhood” (Gray et al., 2003) approach, in which cells surrounding an occupied focal cell (to 1,2…*d* neighbors) may become occupied, depending on their suitability. MigClim instead assumes a probabilistic contagion model that requires parameter estimates for number of propagules, and short- and long-distance-decay rates. Given the lack of empirical data to inform values for those parameters, we prefer a simpler algorithm to explore how connected clusters of suitable cells are across different values of the single parameter *d*. Another factor resulting in these differences is the number of simulation steps used in our approach (200). From a biological perspective, this implies that 200 dispersal events resulting in colonization of suitable cells happened. Although this number may appear excessive, it gives a view of a scenario in which no action is taken to prevent AGH invasion in North America and the species builds to large local populations. For a more conservative view of the expected invasion, one could concentrate in areas with high values on the layers obtained from our simulations.

The areas in North America that our models identified as highly suitable for this hornet overlap broadly with the states where honey production is highest, and species richness of *Bombus* and *Melipona* are highest (Fig. 7). These results give credence to public concerns that, if established, the AGH could pose a serious economic threat to the beekeeping industry in Oregon, northern California, Georgia, Alabama, and Florida. In the United States alone, the European honey bee provides at least $15 billion worth of pollination services and generates between $300 and 500 million in harvestable honey and other products each year (Calderone, 2012). In Mexico, impacts on the honey bee industry are also expected, in tropical areas of the country that have suitable areas for the AGH, particularly in the states of Yucatan, Campeche, and Quintana Roo. Beekeepers in the United States and Mexico may have to adopt mitigation practices to avoid serious losses, such as those developed by Japanese beekeepers including the use of protective screens or traps at the hive entrance that can exclude AGHs based on body size (Matsuura & Sakagami, 1973; Mahdi, Glaiim & Ibrahim, 2008). Potential establishment of the AGH in North America adds an additional layer of environmental and economic stress to a beekeeping industry already suffering from high annual hive mortality rates resulting from the combined effects of pesticides, diseases, and poor nutrition (Goulson et al., 2015).

**Figure 7.**
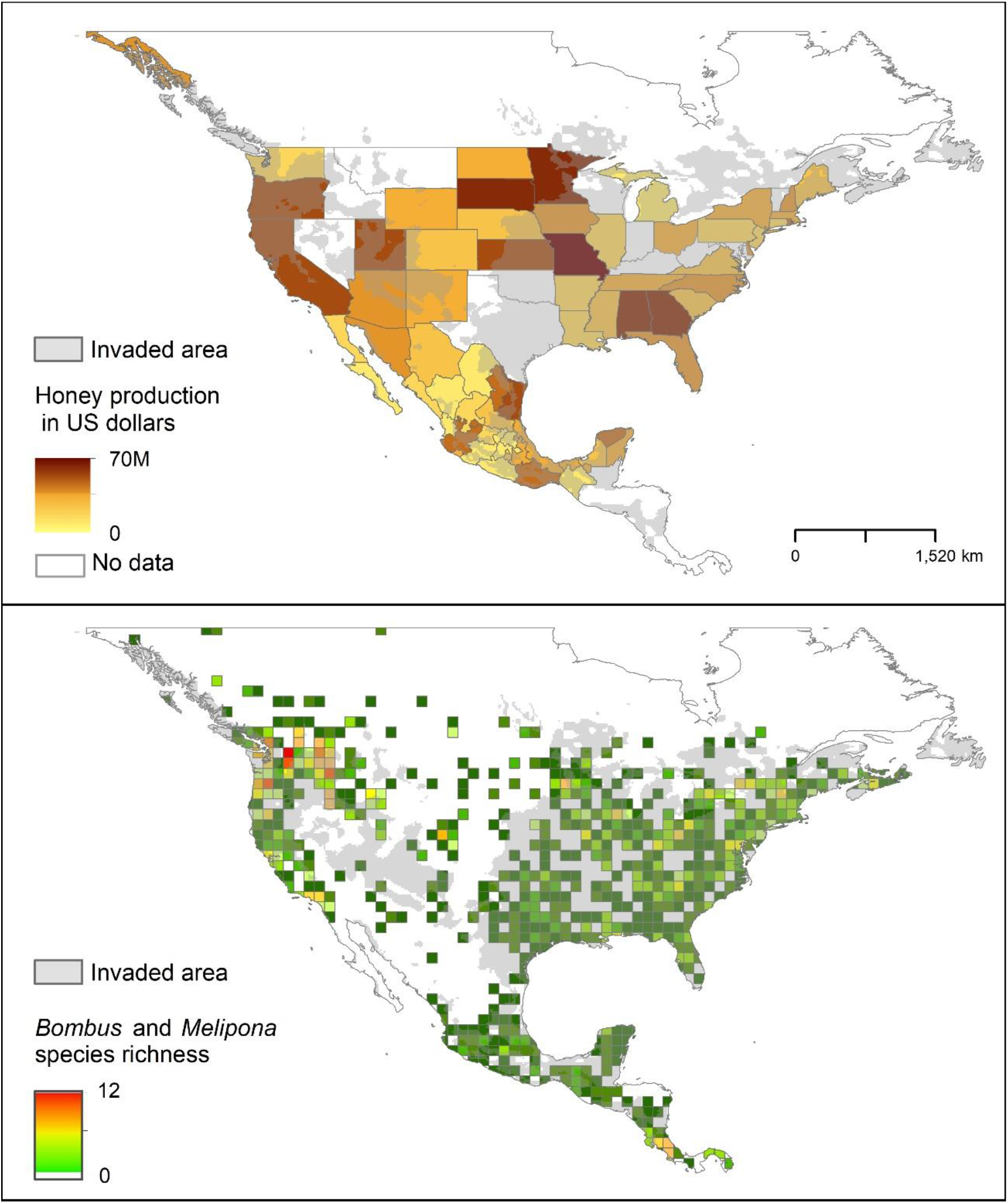
Representation of potential ecological and economic impacts of an invasion of *Vespa mandarinia*. Top panel: honey production (in US dollars) in Mexico and the United States in 2016. Bottom panel: species richness of the genera *Bombus* (bumble bees) and *Melipona* (stingless bees) in North America. The area shaded in gray represents the simulated potential invaded area of *Vespa mandarinia* in North America obtained with the 50 km spatial thinning occurrences and PCs as environmental predictors.

The ecological impact of AGH on the local bee fauna is more challenging to predict than the economic impact on honey production, because it is not clear which native bee species would be particularly targeted by AGH in North America. We explore *Bombus* and *Melipona* species as likely prey candidates of AGH because, among the >4000 bee species occurring in this region (Ascher & Pickering, 2020), these two groups of bees are social, locally abundant, and make annual or perennial colonies (Michener, 2007; Cueva del Castillo, Sanabria-Urbán & Serrano-Meneses, 2015; Viana et al., 2015). Thus, they may represent predictable food sources for the AGH. It is crucial to consider this potential threat because both *Bombus* and *Melipona* bees are important pollinators that have already experienced population losses and local extirpations, reflecting changes in landscape and agricultural intensification (Brown & Albrecht, 2001; Cameron et al., 2011). Furthermore, these species, as well as the European honey bee, lack behavioral responses to prevent predation by the AGH (Matsuura & Sakagami, 1973; McClenaghan et al., 2019), because they have no shared evolutionary history with the AGH, and are thus vulnerable to its predatory and antagonistic behavior. The economic and cultural importance of species of *Melipona* in America is well-documented, particularly in the Yucatan Peninsula in Mexico, where these bees have been traditionally raised for honey and were even considered gods outright in Mayan times (Ayala, Gonzalez & Engel, 2013; Quezada-Euán et al., 2018). It is important to mention, however, that the risk to *Melipona* species may be lower than that to *Bombus* species because entrances to the hives of some species of *Melipona* are narrow, allowing a single bee to pass at a time (Couvillon et al., 2007), unlike the entrances to the hives of honey bees and many bumble bees, which are wider.

The AGH is not the first hornet to invade North America, and species of *Vespa* are well-known for their invasive success and excellent dispersal capacity (Beggs et al., 2011; Monceau, Bonnard & Thiéry, 2014). The solitary giant resin bee, *Challomegachile sculpturalis* is an Asian taxon which was recently introduced in the United States. Only 15 years after its initial detection near Baltimore, Maryland, this species had invaded most of the southeastern United States (Hinojosa-Díaz et al., 2005). Additionally, the European hornet, *Vespa crabro* L., a Eurasian species that was accidentally introduced to North America in the 1800s, occupies a similar invasive range in the United States (Smith et al. 2020). These examples indicate considerable precedent for hornet invasion and establishment in the southeastern United States, but the AGH poses a unique biodiversity risk as a direct predator of bees. Because the Pacific Northwest is consistently predicted as suitable for the AGH, preventing further establishment and spread of recently detected introduced populations near Seattle and Vancouver is essential. If these introduced individuals are not eradicated, they may flourish under the suitable climatic conditions, establishing many more colonies that will be difficult to control. Preventing establishment of the AGH in the Pacific Northwest is especially critical because an established AGH population in the Pacific Northwest would provide a source population for potential long-range dispersers that could use multiple potential invasion routes (Fig. 6) to reach suitable habitat in the eastern United States, facilitating full-scale invasion. In light of this, we recommend official monitoring protocols for the vulnerable Pacific Northwest region, and encourage citizen-science monitoring efforts, which may be the fastest and most effective way to detect potential range expansions.

Although AGH is primarily found in temperate areas in its native range, some of its populations reach subtropical regions like Taiwan (Archer, 2008), which indicates a broad temperature tolerance. This southern part of the species’ native range might explain why our models predicted suitable areas in South America, Africa, and elsewhere (Figure 2S-S7). Although temperature is a critical factor that determines the abundance and distribution of organisms (Sunday, Bates & Dulvy, 2012), factors such as desiccation resistance may be equally important for some species. For example, for ants and some bees, desiccation tolerance is a good predictor of species’ distributions (Bujan, Yanoviak & Kaspari, 2016; Burdine & McCluney, 2019). For example, humidity is important for the regulation of temperature in nests of the European hornet (Klingner et al., 2005) and, in some species of stingless bees, regulation of humidity appears to be more important than regulation of temperature to maintain colony health (Ayton et al., 2016). Unfortunately, heat and desiccation tolerances, factors that might improve predictions of this species’ distributional potential, are unknown for the AGH. In other hornets, subtropical populations tend to have longer population cycles than temperate populations (Archer, 2008), so negative impacts of an AGH invasion may be stronger in tropical or subtropical areas.

In summary, our modeling approach allowed us to recognize how predicted suitable areas can be depending on distinct schemes of data treatment. We showed that this variability can derive from crucial decisions made during the initial steps of ecological niche modeling exercises. These results highlight the importance of such initial decisions, as well as the need to recognize sources of variability. This point is of special importance in predicting the potential for expansion of invasive species, as uncertainty increases when models are transferred to areas where environmental conditions are different. Our analyses and simulations revealed the potential of the AGH to invade large areas in North America and the likely paths of such an invasion. We also showed that predicted suitable areas for the AGH overlap broadly with those where honey production is highest in the United States and Mexico, as well as with species-rich areas for bumble bees and stingless bees. These results bring light to the potential implications of uncontrolled dispersal of the AGH to suitable environments in North America, and highlight the need for rapid eradication actions to mitigate potential biodiversity and economic losses.

## Supporting information

Supplemental files include description of the simulations, and results cited but not included in the text

## Acknowledgments

We would like to thank the members of the KUENM group for their support in the development of this manuscript. We also thank Allan Smith-Pardo for letting us use the photograph of AGH in lateral view (Fig. 1B). ANB would like to thank Secretaría de Educación, Ciencia, Tecnología e Innovación de la Ciudad de México.

## Data Availability

Data is available at KU ScholarWorks http://hdl.handle.net/1808/30602

Code is available at https://github.com/townpeterson/vespa

## Author contributions

CNP, LOO, MEC, LJ, AbA, ATP, and JS conceived and designed the study. CNP, LOO, VHG, MEC, LJ, DAD, AbA, RGCD, ANB, KU, UA, AdA, and JO performed the analysis. CAN, LOO, MEC, and RGCD prepared figures and/or tables. All authors drafted the work or revised critically the manuscript for important content and approved the final version.

## References

APHIS. 2020. New pest response guidelines: *Vespa mandarinia*. U.S. Department of Agriculture, Animal and Plant Health Inspection Service, Plant Protection and Quarantine.

Alkishe A, Cobos ME, Peterson AT, Samy AM. 2020. Recognizing sources of uncertainty in disease vector ecological niche models: an example with the tick *Rhipicephalus sanguineus* sensu lato. Perspectives in Ecology and Conservation. DOI: 10.1016/j.pecon.2020.03.002.

Anderson RP. 2012. Harnessing the world’s biodiversity data: promise and peril in ecological niche modeling of species distributions. Annals of the New York Academy of Sciences 1260:66–80. DOI: 10.1111/j.1749-6632.2011.06440.x.

Anderson RP, Lew D, Peterson AT. 2003. Evaluating predictive models of species’ distributions: criteria for selecting optimal models. Ecological Modelling 162:211–232. DOI: 10.1016/S0304-3800(02)00349-6.

Archer ME. 1995. Taxonomy, distribution and nesting biology of the *Vespa mandarinia* group (Hym., Vespinae). Entomologist’s Monthly Magazine 131:47–53.

Archer ME. 2008. Taxonomy, distribution and nesting biology of species of the genera Provespa Ashmead and *Vespa* Linneaus (Hymenoptera, Vespidae). Entomologist’s Monthly Magazine 144:69–101.

Arita HT, Christen JA, Rodríguez P, Soberón J. 2008. Species diversity and distribution in presence-absence matrices: mathematical relationships and biological implications. American Naturalist 172:519–532. DOI: 10.1086/590954.

Ascher JS, Pickering J. 2020. Discover life bee species guide and world checklist (Hymenoptera: Apoidea: *Anthophila*). Available at https://www.discoverlife.org/mp/20q?guide=Apoidea_species (accessed July 1, 2020).

Ayala R, Gonzalez VH, Engel MS. 2013. Mexican stingless bees (Hymenoptera: Apidae): diversity, distribution, and indigenous knowledge. In: Vit P, Pedro SRM, Roubik D eds. Pot-Honey: A Legacy of Stingless Bees. New York, NY: Springer, 135–152. DOI: 10.1007/978-1-4614-4960-7_9.

Ayton S, Tomlinson S, Phillips RD, Dixon KW, Withers PC. 2016. Phenophysiological variation of a bee that regulates hive humidity, but not hive temperature. Journal of Experimental Biology 219:1552–1562. DOI: 10.1242/jeb.137588.

Barve N, Barve V, Jiménez-Valverde A, Lira-Noriega A, Maher SP, Peterson AT, Soberón J, Villalobos F. 2011. The crucial role of the accessible area in ecological niche modeling and species distribution modeling. Ecological Modelling 222:1810–1819. DOI: 10.1016/j.ecolmodel.2011.02.011.

Beggs JR, Brockerhoff EG, Corley JC, Kenis M, Masciocchi M, Muller F, Rome Q, Villemant C. 2011. Ecological effects and management of invasive alien Vespidae. BioControl 56:505–526. DOI: 10.1007/s10526-011-9389-z.

Bérubé C. 2020. Giant alien insect invasion averted Canadian beekeepers thwart apicultural disaster (… or at least the zorn-bee apocalypse). American Bee Journal:209–214.

Bivand R, Keitt T, Rowlingson B, Pebesma E, Sumner M, Hijmans R, Rouault E, Warmerdam F, Ooms J, Rundel C. 2020a. rgdal: Bindings for the “geospatial” data abstraction library. https://cran.r-project.org/web/packages/rgdal/index.html.

Bivand R, Rundel C, Pebesma E, Stuetz R, Hufthammer KO, Giraudoux P, Davis M, Santilli S. 2020b. rgeos: Interface to geometry engine - open source (‘GEOS’). https://cran.r-project.org/web/packages/rgeos/index.html.

Brown JC, Albrecht C. 2001. The effect of tropical deforestation on stingless bees of the genus *Melipona* (Insecta: Hymenoptera: Apidae: Meliponini) in central Rondonia, Brazil. Journal of Biogeography 28:623–634. DOI: 10.1046/j.1365-2699.2001.00583.x.

Bujan J, Yanoviak SP, Kaspari M. 2016. Desiccation resistance in tropical insects: causes and mechanisms underlying variability in a Panama ant community. Ecology and Evolution 6:6282–6291. DOI: 10.1002/ece3.2355.

Burdine JD, McCluney KE. 2019. Differential sensitivity of bees to urbanization-driven changes in body temperature and water content. Scientific Reports 9:1643. DOI: 10.1038/s41598-018-38338-0.

Cameron SA, Lozier JD, Strange JP, Koch JB, Cordes N, Solter LF, Griswold TL. 2011. Patterns of widespread decline in North American bumble bees. Proceedings of the National Academy of Sciences USA 108:662–667. DOI: 10.1073/pnas.1014743108.

Cobos ME, Jiménez L, Nuñez-Penichet C, Romero-Alvarez D, Simões M. 2018. Sample data and training modules for cleaning biodiversity information. Biodiversity Informatics 13:49–50. DOI: 10.17161/bi.v13i0.7600.

Cobos ME, Osorio-Olvera L, Soberón J, Peterson AT, Barve V, Barve N. 2020. ellipsenm: Ecological niche characterizations using ellipsoids. https://github.com/marlonecobos/ellipsenm.

Cobos ME, Peterson AT, Barve N, Osorio-Olvera L. 2019a. kuenm: an R package for detailed development of ecological niche models using Maxent. PeerJ 7:e6281. DOI: 10.7717/peerj.6281.

Cobos ME, Peterson AT, Osorio-Olvera L, Jiménez-García D. 2019b. An exhaustive analysis of heuristic methods for variable selection in ecological niche modeling and species distribution modeling. Ecological Informatics 53:100983. DOI: 10.1016/j.ecoinf.2019.100983.

Couvillon MJ, Wenseleers T, Imperatriz-Fonseca VL, Nogueira-Neto P, Ratnieks FLW. 2007. Comparative study in stingless bees (Meliponini) demonstrates that nest entrance size predicts traffic and defensivity. Journal of Evolutionary Biology 21:194–201. DOI: 10.1111/j.1420-9101.2007.01457.x.

Cueva del Castillo R, Sanabria-Urbán S, Serrano-Meneses MA. 2015. Trade-offs in the evolution of bumblebee colony and body size: a comparative analysis. Ecology and Evolution 5:3914–3926. DOI: 10.1002/ece3.1659.

Dueñas M-A, Ruffhead HJ, Wakefield NH, Roberts PD, Hemming DJ, Diaz-Soltero H. 2018. The role played by invasive species in interactions with endangered and threatened species in the United States: a systematic review. Biodiversity and Conservation 27:3171–3183. DOI: 10.1007/s10531-018-1595-x.

Engler R, Hordijk W, Guisan A. 2012. The MIGCLIM R package – seamless integration of dispersal constraints into projections of species distribution models. Ecography 35:872–878. DOI: 10.1111/j.1600-0587.2012.07608.x.

Escobar LE, Lira-Noriega A, Medina-Vogel G, Peterson AT. 2014. Potential for spread of the white-nose fungus (*Pseudogymnoascus destructans*) in the Americas: use of Maxent and NicheA to assure strict model transference. Geospatial Health 9:221–229. DOI: 10.4081/gh.2014.19.

Fujiwara A, Sasaki M, Washitani I. 2016. A scientific note on hive entrance smearing in Japanese *Apis cerana* induced by pre-mass attack scouting by the Asian giant hornet Vespa mandarinia. Apidologie 47:789–791. DOI: 10.1007/s13592-016-0432-z.

GBIF.org (07 May 2020) GBIF occurrence download. https://doi.org/10.15468/dl.kzcgc2.

Gray L, New A, Science K, Wolfram S. 2003. A mathematician looks at Wolfram’s new kind of science. Notices of the American Mathematical Society 50 (2) (2003) 200–211, URL http://www.ams.org/notices/200302/fea-gray.pdf. URL http://www.ams.org/notices/200302/fea-gray.pdf 50:200–211.

Hijmans RJ, Etten J van, Sumner M, Cheng J, Bevan A, Bivand R, Busetto L, Canty M, Forrest D, Ghosh A, Golicher D, Gray J, Greenberg JA, Hiemstra P, Hingee K, Geosciences I for MA, Karney C, Mattiuzzi M, Mosher S, Nowosad J, Pebesma E, Lamigueiro OP, Racine EB, Rowlingson B, Shortridge A, Venables B, Wueest R. 2020. raster: Geographic data analysis and modeling. https://cran.r-project.org/web/packages/raster/index.html.

Hinojosa-Díaz IA, Yáñez-Ordóñez O, Chen G, Peterson AT, Engel MS. 2005. The North American invasion of the giant resin bee (Hymenoptera: Megachilidae). Journal of Hymenoptera Research 14:69–77.

Kastberger G, Schmelzer E, Kranner I. 2008. Social waves in Giant Honeybees repel hornets. PLoS ONE 3:e3141. DOI: 10.1371/journal.pone.0003141.

Klingner R, Richter K, Schmolz E, Keller B. 2005. The role of moisture in the nest thermoregulation of social wasps. Naturwissenschaften 92:427–430. DOI: 10.1007/s00114-005-0012-y.

Li X-D, Liu Z, Zhai Y, Zhao M, Shen H-Y, Li Y, Zhang B, Liu T. 2015. Acute interstitial nephritis following multiple Asian Giant Hornet stings. American Journal of Case Reports 16:371–373. DOI: 10.12659/AJCR.893734.

Matsuura M. 1988. Ecological study on Vespine wasps (Hymenoptera: Vespidae) attacking honeybee colonies: 1. Seasonal changes in the frequency of visits to apiaries by Vespine wasps and damage inflicted, especially in the absence of artificial protection. Applied Entomology and Zoology 23:428–440.

Matsuura M, Sakagami SF. 1973. A bionomic sketch of the Giant Hornet, *Vespa mandarinia*, a serious pest for Japanese apiculture. Journal of the Faculty of Science, Hokkaido University: Series 6. Zoology 19:125–162.

McClenaghan B, Schlaf M, Geddes M, Mazza J, Pitman G, McCallum K, Rawluk S, Hand K, Otis GW. 2019. Behavioral responses of honey bees, *Apis cerana* and *Apis mellifera*, to Vespa mandarinia marking and alarm pheromones. Journal of Apicultural Research 58:141–148. DOI: 10.1080/00218839.2018.1494917.

Michener CD. 2000. The Bees of the World. Baltimore: Johns Hopkins University Press.

Michener CD. 2007. The Bees of the World. Baltimore: Johns Hopkins University Press.

Monceau K, Bonnard O, Thiéry D. 2014. *Vespa velutina*: a new invasive predator of honeybees in Europe. Journal of Pest Science 87:1–16. DOI: 10.1007/s10340-013-0537-3.

Nuñez-Penichet C, Cobos ME, Peterson AT, Barve N, Barve V, Gueta T, Soberón J. 2020. biosurvey: Tools for biological survey planning. https://github.com/claununez/biosurvey.

Ono M, Igarashi T, Ohno E, Sasaki M. 1995. Unusual thermal defense by a honeybee against mass attack by hornets. Nature 377:334–336. DOI: 10.1038/377334a0.

Osorio-Olvera L, Soberón J. 2020. bam: Species distribution models in the light of the BAM theory. https://github.com/luismurao/bam.

Osorio-Olvera L, Soberón J, Barve V, Barve N, Falconi M. 2020. ntbox: From getting biodiversity data to evaluating species distribution models in a friendly GUI environment. https://github.com/luismurao/ntbox.

Owens HL, Campbell LP, Dornak LL, Saupe EE, Barve N, Soberón J, Ingenloff K, Lira-Noriega A, Hensz CM, Myers CE, Peterson AT. 2013. Constraints on interpretation of ecological niche models by limited environmental ranges on calibration areas. Ecological Modelling 263:10–18. DOI: 10.1016/j.ecolmodel.2013.04.011.

Peterson AT, Papeş M, Soberón J. 2008. Rethinking receiver operating characteristic analysis applications in ecological niche modeling. Ecological Modelling 213:63–72. DOI: 10.1016/j.ecolmodel.2007.11.008.

Phillips SJ, Anderson RP, Dudík M, Schapire RE, Blair ME. 2017. Opening the black box: an open-source release of Maxent. Ecography 40:887–893. DOI: 10.1111/ecog.03049.

Phillips SJ, Anderson RP, Schapire RE. 2006. Maximum entropy modeling of species geographic distributions. Ecological Modelling 190:231–259. DOI: 10.1016/j.ecolmodel.2005.03.026.

Pimentel D, Zuniga R, Morrison D. 2005. Update on the environmental and economic costs associated with alien-invasive species in the United States. Ecological Economics 52:273–288. DOI: 10.1016/j.ecolecon.2004.10.002.

Pyšek P, Richardson DM. 2010. Invasive species, environmental change and management, and health. Annual Review of Environment and Resources 35:25–55. DOI: 10.1146/annurev-environ-033009-095548.

Quezada-Euán JJG, Nates-Parra G, Maués MM, Roubik DW, Imperatriz-Fonseca VL. 2018. The economic and cultural values of stingless bees (Hymenoptera: Meliponini) among ethnic groups of tropical America. Sociobiology 65:534–557. DOI: 10.13102/sociobiology.v65i4.3447.

R Core Team. 2019. R: A language and environment for statistical computing. Vienna, Austria: R Foundation for Statistical Computing.

Schmidt JO, Yamane S, Matsuura M, Starr CK. 1986. Hornet venoms: lethalities and lethal capacities. Toxicon 24:950–954. DOI: 10.1016/0041-0101(86)90096-6.

Simões M, Romero-Alvarez D, Nuñez-Penichet C, Jiménez L, Cobos ME. 2020. General theory and good practices in ecological niche modeling: a basic guide. Biodiversity Informatics 15:67–68.

Smith-Pardo AH, Carpenter JM, Kimsey L. 2020. The diversity of hornets in the genus Vespa (Hymenoptera: Vespidae; Vespinae), their importance and interceptions in the United States. Insect Systematics and Diversity 4:1–27.

Sugahara M, Nishimura Y, Sakamoto F. 2012. Differences in heat sensitivity between Japanese honeybees and hornets under high carbon dioxide and humidity conditions inside bee balls. Zoological Science 29:30–36. DOI: 10.2108/zsj.29.30.

Sunday JM, Bates AE, Dulvy NK. 2012. Thermal tolerance and the global redistribution of animals. Nature Climate Change 2:686–690. DOI: 10.1038/nclimate1539.

Vega GC, Pertierra LR, Olalla-Tárraga MÁ. 2018. MERRAclim, a high-resolution global dataset of remotely sensed bioclimatic variables for ecological modelling. Scientific Data 4:170078. DOI: 10.1038/sdata.2017.78.

Viana JL, Sousa H de AC, Alves RM de O, Pereira DG, Silva Jr. JC, Paixão JF da, Waldschmidt AM, Viana JL, Sousa H de AC, Alves RM de O, Pereira DG, Silva Jr. JC, Paixão JF da, Waldschmidt AM. 2015. Bionomics of *Melipona mondury* Smith 1863 (Hymenoptera: Apidae, Meliponini) in relation to its nesting behavior. Biota Neotropica 15. DOI: 10.1590/1676-06032015009714.

Vilà M, Espinar JL, Hejda M, Hulme PE, Jarošík V, Maron JL, Pergl J, Schaffner U, Sun Y, Pyšek P. 2011. Ecological impacts of invasive alien plants: a meta-analysis of their effects on species, communities and ecosystems. Ecology Letters 14:702–708. DOI: 10.1111/j.1461-0248.2011.01628.x.

Warren DL, Seifert SN. 2011. Ecological niche modeling in Maxent: the importance of model complexity and the performance of model selection criteria. Ecological Applications 21:335–342. DOI: 10.1890/10-1171.1.

Wilcove DS, Rothstein D, Dubow J, Phillips A, Losos E. 1998. Quantifying threats to imperiled species in the United States. BioScience 48:607–615. DOI: 10.2307/1313420.

Yanagawa Y, Morita K, Sugiura T, Okada Y. 2007. Cutaneous hemorrhage or necrosis findings after *Vespa mandarinia* (wasp) stings may predict the occurrence of multiple organ injury: a case report and review of literature. Clinical Toxicology 45:803–807. DOI: 10.1080/15563650701664871.

Zhu G, Illan JG, Looney C, Crowder DW. 2020. Assessing the ecological niche and invasion potential of the Asian giant hornet. bioRxiv. DOI: 10.1101/2020.05.25.115311.

Zurell D, Franklin J, König C, Bouchet PJ, Dormann CF, Elith J, Fandos G, Feng X, Guillera-Arroita G, Guisan A, Lahoz-Monfort JJ, Leitão PJ, Park DS, Peterson AT, Rapacciuolo G, Schmatz DR, Schröder B, Serra-Diaz JM, Thuiller W, Yates KL, Zimmermann NE, Merow C. 2020. A standard protocol for reporting species distribution models. Ecography. DOI: 10.1111/ecog.04960.

