## Supplemental files include description of the simulations, and results cited but not included in the text for "Geographic potential of the world’s largest hornet, *Vespa mandarinia* Smith (Hymenoptera: Vespidae), worldwide and particularly in North America"

**Supporting Information**

Table S1. Sets of environmental predictors obtained with all combinations of two or more raw variables (BIO3: isothermality, BIO5: maximum temperature of warmest month, BIO6: minimum temperature of coldest month, BIO7: temperature annual range, BIO13: specific humidity of most humid month, BIO14: specific humidity of least humid month).

| Set name | BIO3 | BIO5 | BIO6 | BIO7 | BIO13 | BIO14 |
|---|---|---|---|---|---|---|
| Set 1 | | | | | X | X |
| Set 2 | X | | | | X | |
| Set 3 | | X | | | X | |
| Set 4 | | | X | | X | |
| Set 5 | | | | X | X | |
| Set 6 | | | X | | | X |
| Set 7 | | X | | | | X |
| Set 8 | | | X | | | X |
| Set 9 | | | | X | | X |
| Set 10 | X | X | | | | |
| Set 11 | X | | X | | | |
| Set 12 | X | | | X | | |
| Set 13 | | X | X | | | |
| Set 14 | | X | | X | | |
| Set 15 | | | X | X | | |
| Set 16 | X | | | | X | X |
| Set 17 | | X | | | X | X |
| Set 18 | | | X | | X | X |
| Set 19 | | | | X | X | X |
| Set 20 | X | X | | | X | |
| Set 21 | X | | X | | X | |
| Set 22 | X | | | X | X | |
| Set 23 | | X | X | | X | |
| Set 24 | | X | | X | X | |
| Set 25 | | | X | X | X | |
| Set 26 | X | X | | | | X |
| Set 27 | X | | X | | | X |
| Set 28 | X | | | X | | X |
| Set 29 | | X | X | | | X |

| | | | | | | |
|---|---|---|---|---|---|---|
| Set 30 | | X | | X | | X |
| Set 31 | | | X | X | | X |
| Set 32 | X | X | X | | | |
| Set 33 | X | X | | X | | |
| Set 34 | X | | X | X | | |
| Set 35 | | X | X | X | | |
| Set 36 | X | X | | | X | X |
| Set 37 | X | | X | | X | X |
| Set 38 | X | | | X | X | X |
| Set 39 | | X | X | | X | X |
| Set 40 | | X | | X | X | X |
| Set 41 | | | X | X | X | X |
| Set 42 | X | X | X | | X | |
| Set 43 | X | X | | X | X | |
| Set 44 | X | | X | X | X | |
| Set 45 | | X | X | X | X | |
| Set 46 | X | X | X | | | X |
| Set 47 | X | X | | X | | X |
| Set 48 | X | | X | X | | X |
| Set 49 | | X | X | X | | X |
| Set 50 | X | X | X | X | | |
| Set 51 | X | X | X | | X | X |
| Set 52 | X | X | | X | X | X |
| Set 53 | X | | X | X | X | X |
| Set 54 | | X | X | X | X | X |
| Set 55 | X | X | X | X | X | |
| Set 56 | X | X | X | X | | X |
| Set 57 | X | X | X | X | X | X |

Table S2. Sets of environmental predictors obtained with all combinations of two or more principal components (PCs).

| Set name | PC1 | PC2 | PC3 | PC4 |
|---|---|---|---|---|
| Set 1 | X | X | | |
| Set 2 | X | | X | |
| Set 3 | X | | | X |
| Set 4 | | X | X | |
| Set 5 | | X | | X |
| Set 6 | | | X | X |
| Set 7 | X | X | X | |
| Set 8 | X | X | | X |
| Set 9 | X | | X | X |
| Set 10 | | X | X | X |
| Set 11 | X | X | X | X |

Table S3. Prevalence of suitable areas obtained for *Vespa mandarinia* in North America, before and after trimming the models with their MOP.

| MOP trimming | Model scheme | Total area (km$^2$) | Area predicted as suitable (km$^2$) | Prevalence |
|---|---|---|---|---|
| No trimming | Raw variables and distance thinned occurrences | 20965151.288 | 3839474.074 | 0.183 |
| | PCs and distance thinned occurrences | 20965151.288 | 6453843.348 | 0.308 |
| | Raw variables and country-density thinned occurrences | 20965151.288 | 3774173.917 | 0.180 |
| | PCs and country-density thinned | 20965151.288 | 6863510.138 | 0.327 |
| After trimming | Raw variables and distance thinned occurrences | 20965151.288 | 3580694.551 | 0.171 |
| | PCs and distance thinned occurrences | 20965151.288 | 5201290.428 | 0.248 |
| | Raw variables and country-density thinned occurrences | 20965151.288 | 3428581.505 | 0.164 |
| | PCs and country-density thinned | 20965151.288 | 5006070.31 | 0.239 |

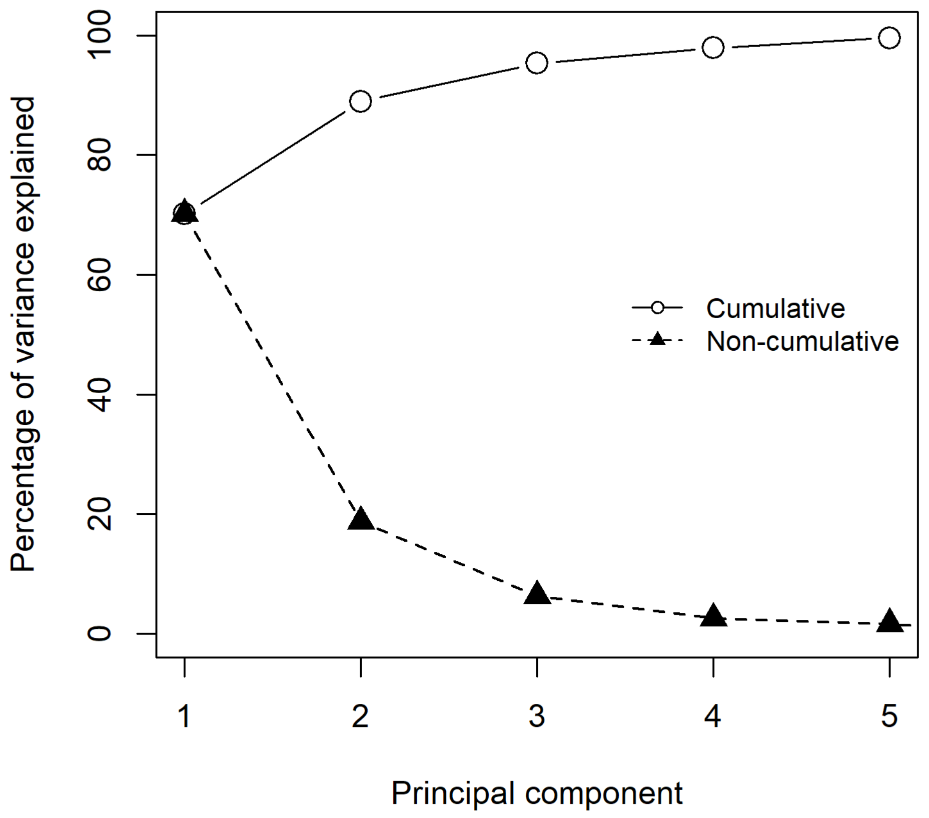


Figure S1. Percentage of variance explained for the first 5 principal components.

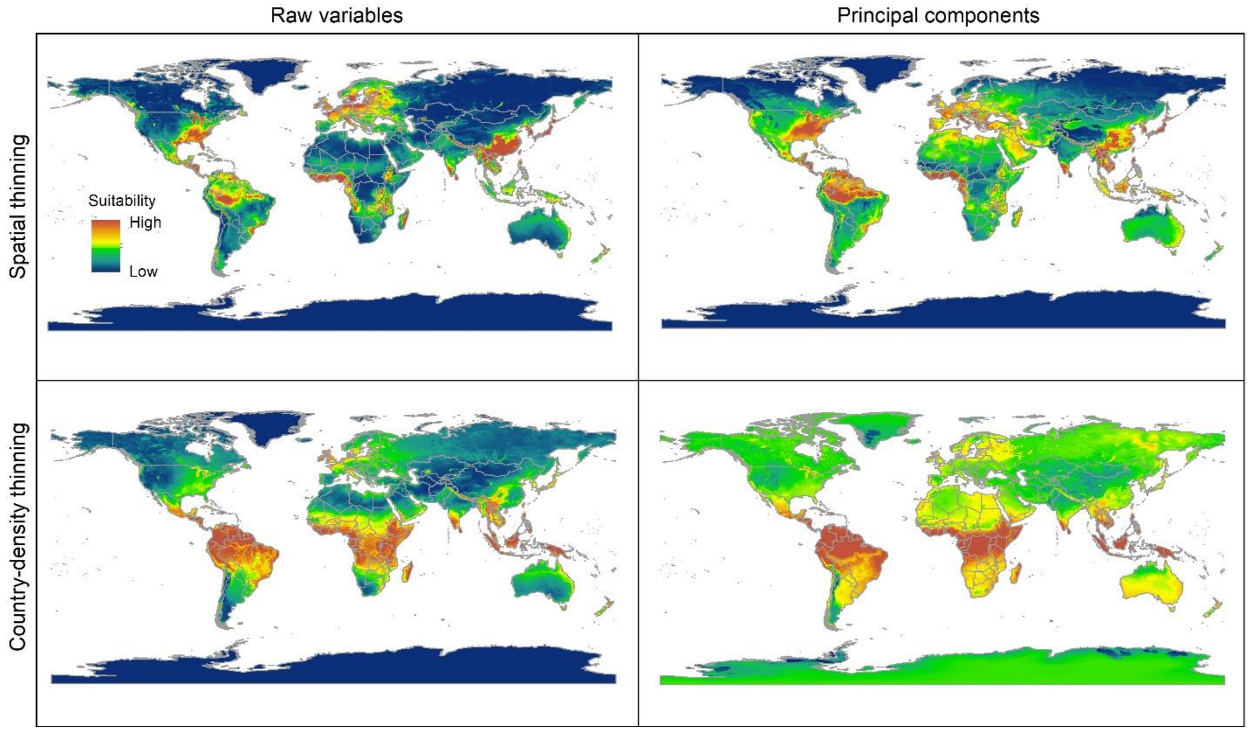


Figure S2. Median of potentially suitable areas for *Vespa mandarinia* predicted with free extrapolation for different calibration schemes projected to the world.

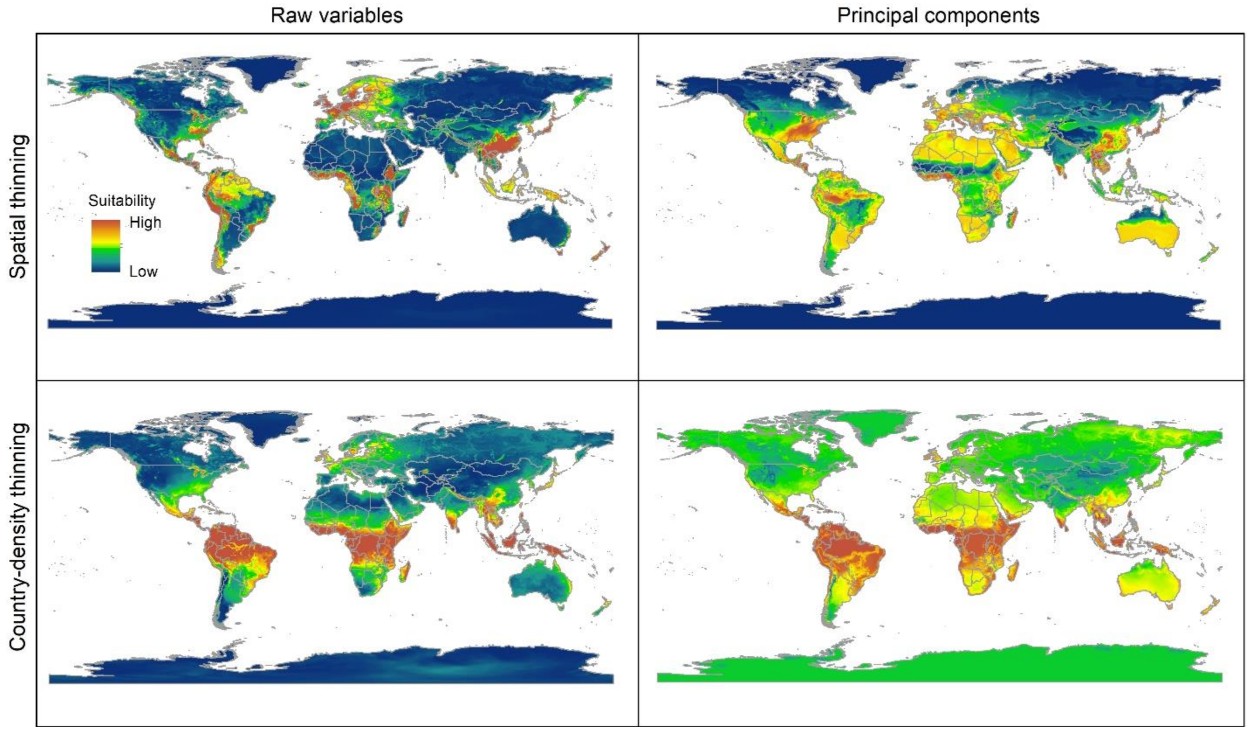


Figure S3. Median of potentially suitable areas for *Vespa mandarinia* predicted with extrapolation and clamping for different calibration schemes projected to the world.

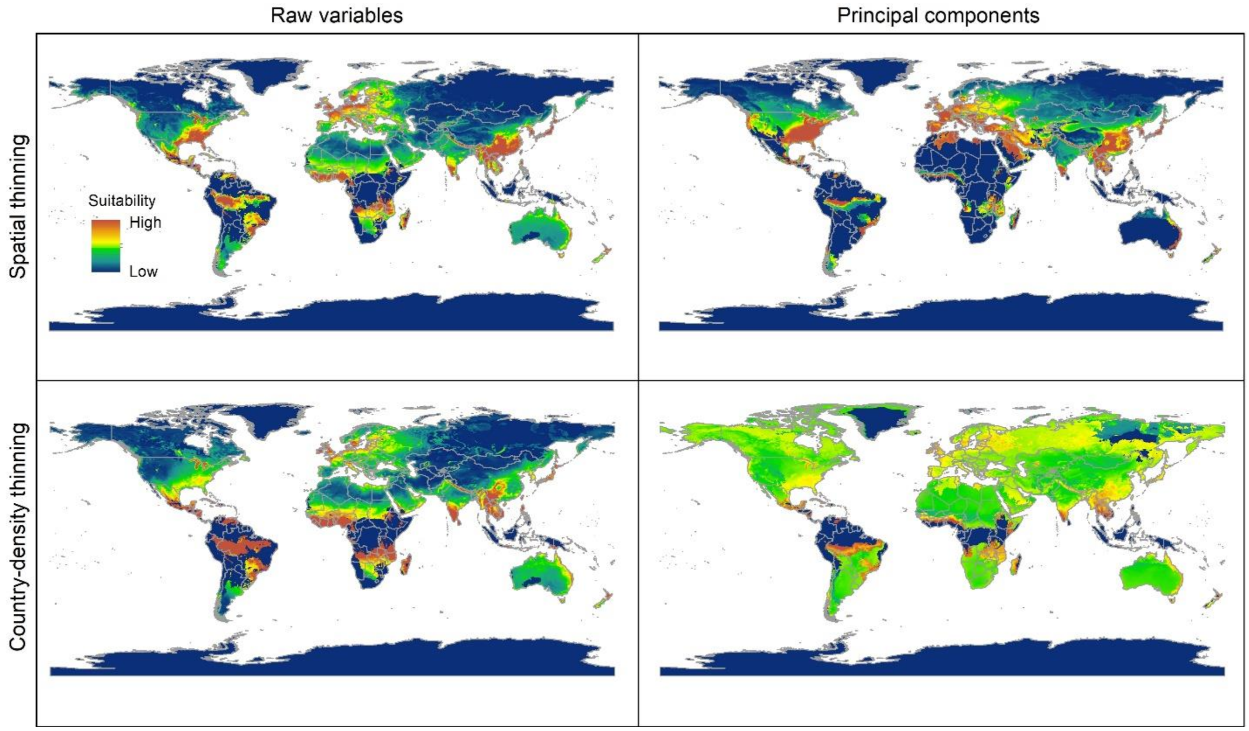


Figure S4. Median of potentially suitable areas for *Vespa mandarinia* predicted with no extrapolation for different calibration schemes projected to the world.

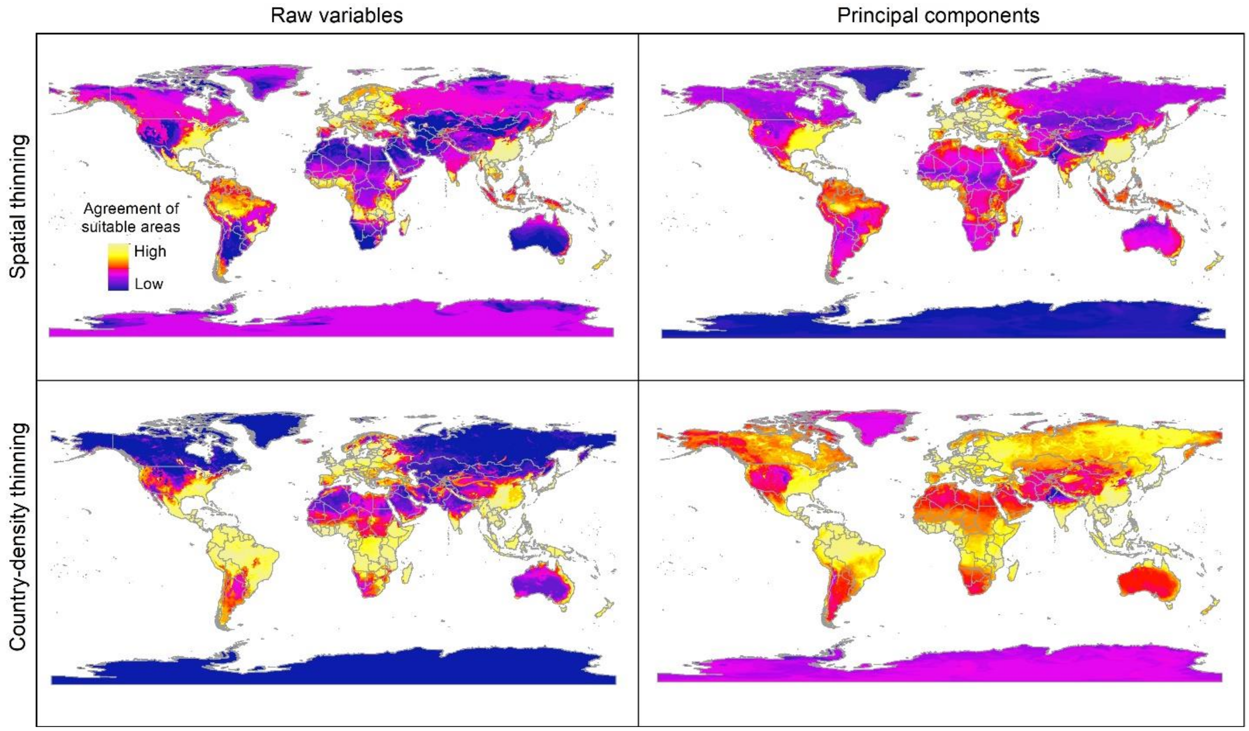


Figure S5. Sum of all suitable areas for *Vespa mandarinia* derived from binarizing (using a 5% threshold) each replicate of selected models (done with free extrapolation) that predicted the known invaded localities of this hornet.

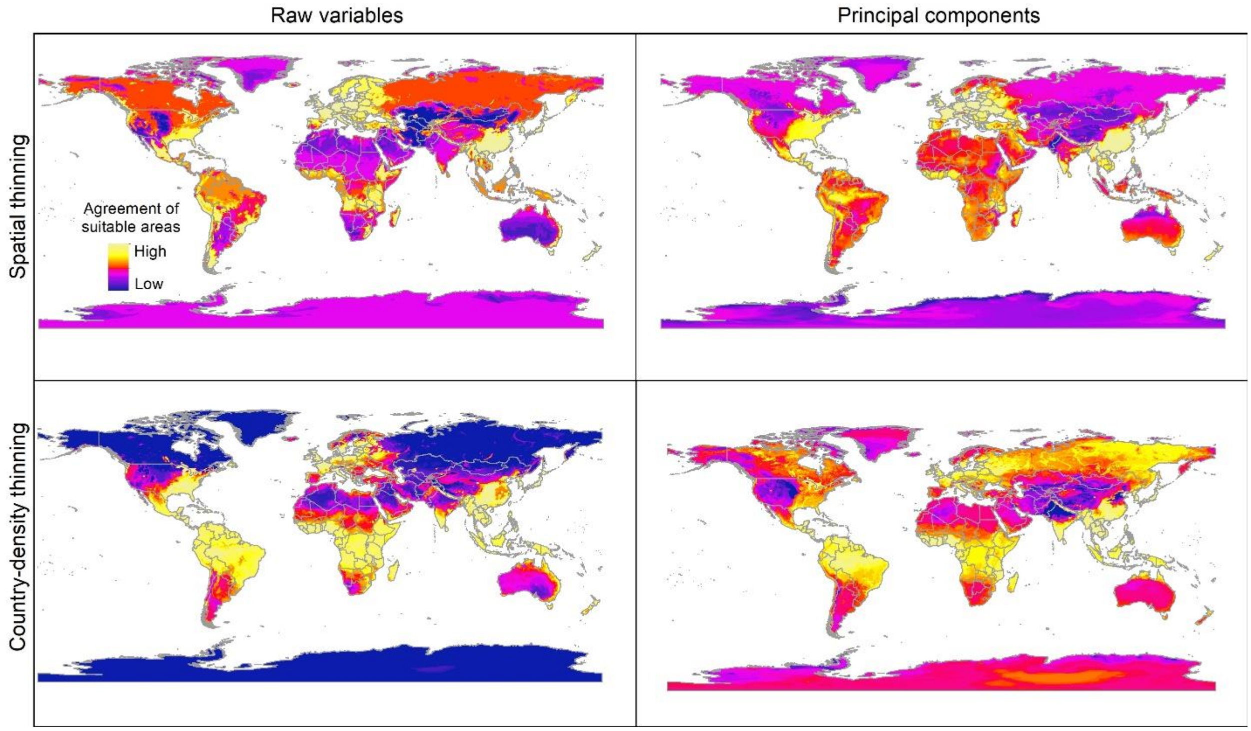


Figure S6. Sum of all suitable areas for *Vespa mandarinia* in North America derived from binarizing (using a 5% threshold) each replicate of selected models (done with extrapolation and clamping) that predicted the known invaded localities of this hornet.

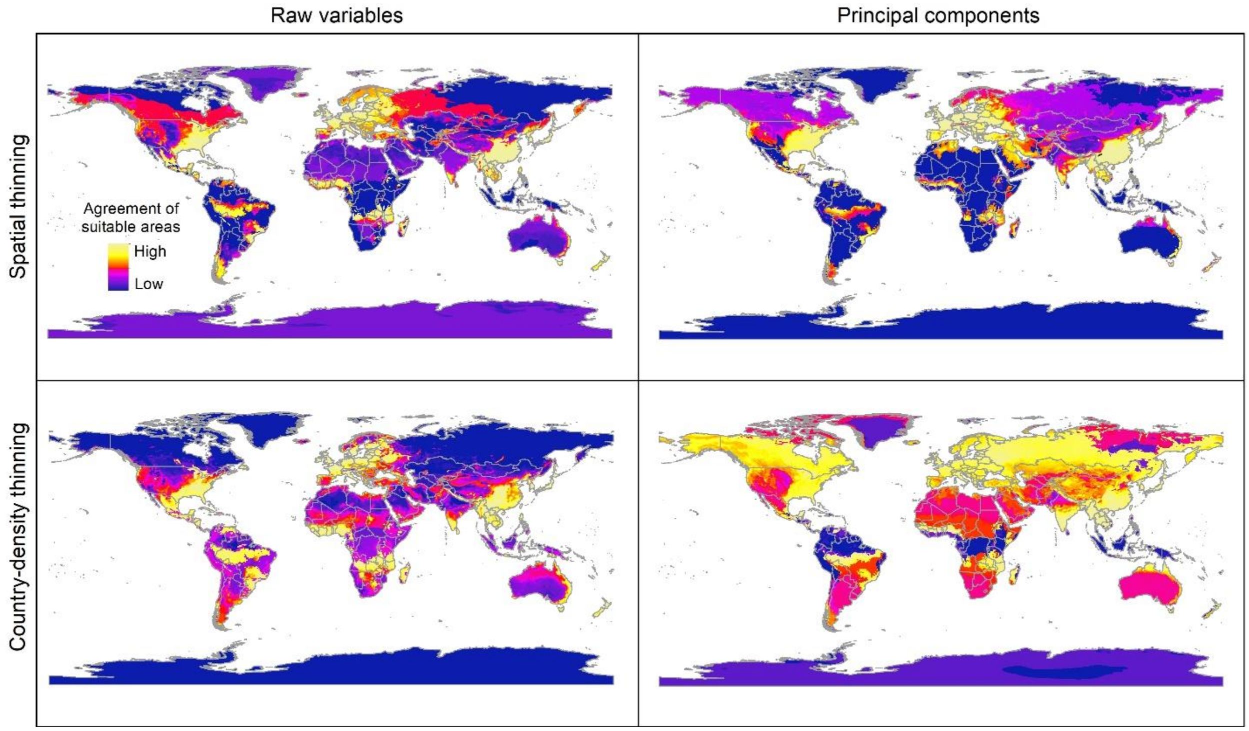


Figure S7. Sum of all suitable areas for *Vespa mandarinia* in North America derived from binarizing (using a 5% threshold) each replicate of selected models (done with no extrapolation) that predicted the known invaded localities of this hornet.

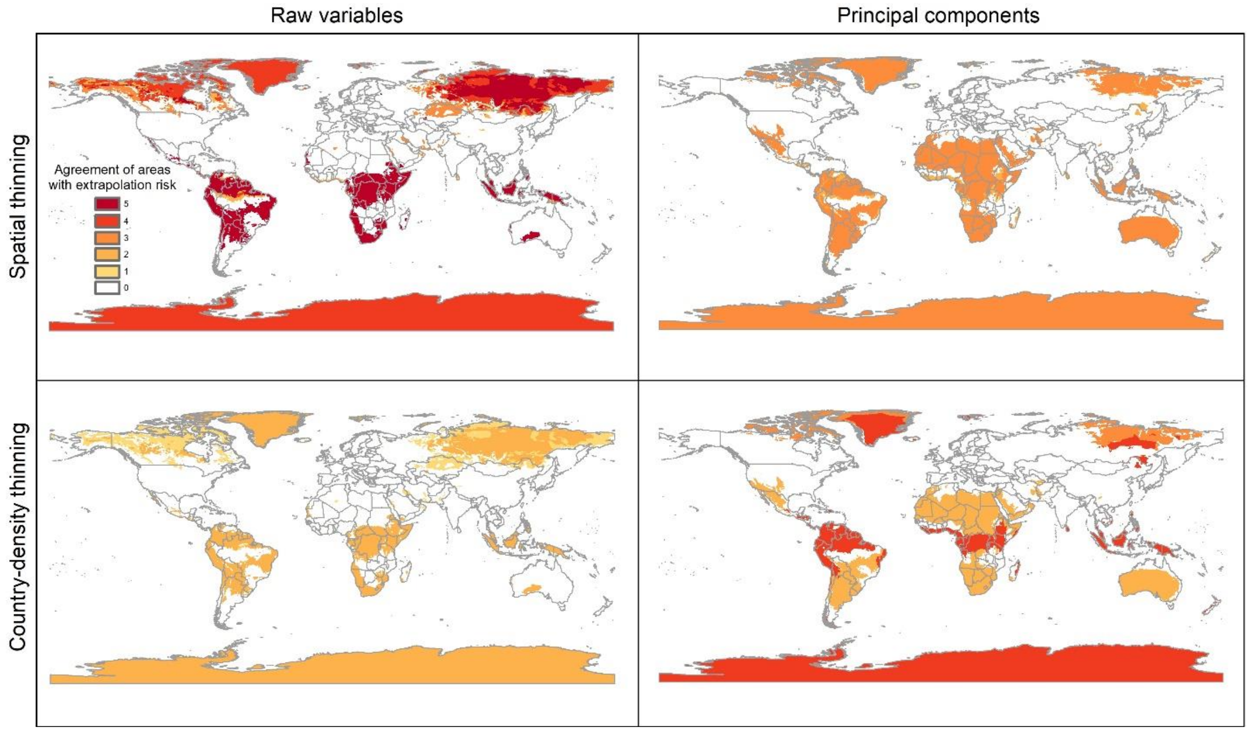


Figure S8. Agreement of areas with extrapolation risk of models of *Vespa mandarinia* worldwide.

**Simulations of the likely invasion dynamics of *Vespa mandarinia***

Using the known invaded localities of the AGH in North America as a starting point, we performed simulations of the potential invasion dynamics of this species. All simulations were done in R using the bam package (https://github.com/luismurao/bam), which is based on a cellular automaton discrete model and simulates how species distributions change as a function of the dispersal ability of the species, given an ecological niche model. Here, we describe the steps we followed to create the different invasion scenarios.

To obtain the occupied area at time $t+1$, for each of the schemes considered to obtain ecological niche models for the species, the algorithm calculates the product of three matrices: (i) the suitability matrix for the species (also called the binary map), (ii) the connectivity matrix that defines the connected patches where the species has the ability to reach in one unit of time, and (iii) the matrix of occupied area at time T. Besides considering the four schemes used to obtain the ecological niche models, we created different invasion scenarios by varying (1) the dispersal parameter, *d*, that determines how far the species is able to move in one unit of time; and (2) the suitability threshold used to create the binary maps. The box below describes the algorithm we used to generate simulations for all scenarios.

Notice that we are using ten different values of dispersal and ten different suitability thresholds (steps 1 and 2, respectively). Therefore, after finishing Step 4, we end up with 100 connectivity maps which will be then added together in step 5 to generate the final distribution map under each scheme. Given that the connectivity maps only indicate whether a cell is occupied or not (i.e., they contain only zeros and ones), adding all the connectivity maps together results in cell values from 0 to 100, where 0 indicates that the corresponding cell was never reached by the species, and a value of 100 indicates that 100 of the times the species was able to occupy that cell. Finally, these values can be also used to visualize the dynamics of the invasion through time. We created an animated map by varying the dispersal parameter, d, where we can see the order in which the different cells are reached by the species, indicating the route of potential invasion of *V. mandarinia*.

**Simulation steps**

1. Generate connectivity matrices, $\mathbf{C}_d$, for each dispersal value $d \in D = \{1, 2, \ldots, 8, 10, 12\}$.

2. Get the suitability thresholds that correspond to the 3% and 10% of the presence points used to obtain an ecological niche model. Let $u_1$ and $u_{10}$ be the suitability thresholds that correspond to 3% and 10% of the presence points, respectively.

3. Create a partition of size 10 of the interval $[u_1, u_{10}]$: $T = [u_1, u_2, \ldots, u_9, u_{10}]$, where $u_1 < u_i < u_{10}, i = 2, \ldots, 9$.

4. Repeat the following steps for each $u_i \in T$ and each $d \in D$:

   (a) Create a binary map, $\mathbf{A}_i$, with the threshold $u_i$.

   (b) Iterate equation (1) for the time steps $t = 1, 2, \ldots, 200$ using the AGH presence point of North America as the initial condition of the distribution ($t = 0$):

   $$\mathbf{G}_{i,d}(t+1) = \mathbf{A}_i \mathbf{C}_d \mathbf{G}(t) \tag{1}$$

   (c) Save the matrix $\mathbf{G}_{i,d}(200)$ which represents the distribution of the species at time $t = 200$.

5. Add the matrices, $\mathbf{G}_{i,d}(200)$, for all $u_i \in T$ and each $d \in D$. Save the resulting matrix which contains the final distribution map of the species for a given ecological niche model.

6. Repeat steps $1 - 5$ for each ecological niche model scheme considered in the study.

### Geographic potential of the world's largest hornet, *Vespa mandarinia* Smith (Hymenoptera: Vespidae), worldwide and particularly in North America

#### – ODMAP Protocol –

Claudia Nuñez-Penichet, Luis Osorio-Olvera, Victor Gonzalez, Marlon E. Cobos, Laura Jiménez, Devon A. DeRaad, Abdelghafar Alkishe, Rusby G. Contreras-Díaz, Angela Nava-Bolaños, Kaera Utsumi, Uzma Ashraf, Adeola Adeboje, A. Townsend Peterson, Jorge Soberón

2020-08-10

---

#### Overview

##### Authorship

Contact:

<Study link>

##### Model objective

Model objective: Inference and explanation

##### Focal Taxon

Focal Taxon: *Vespa mandarinia*

##### Location

Location: North America, Southeast Asia, world

##### Scale of Analysis

Spatial extent: -180, 180, -90, 83.623596 (xmin, xmax, ymin, ymax)

Spatial resolution: 10' resolution (~18 km at the Equator)

Temporal extent: 1990-2020

Temporal resolution: N/A

Boundary: natural

#### Biodiversity data

Observation type: citizen science, field survey, GPS tracking, standardised monitoring data

Response data type: presence-only

#### Predictors

Predictor types: climatic

#### Hypotheses

Hypotheses: Dispersal simulations based on invasive records and areas detected to be suitable using ecological niche models allow detection of patterns of potential invasion of *V. mandarinia* in North America.

#### Assumptions

Model assumptions: After data cleaning and thinning, species data are free of bias. Predictors are free of bias after initial steps of selection.

#### Algorithms

Modelling techniques: Maxent

Model complexity: A process of variable selection (based on correlation and biological relevance) and principal component analysis was used to control the complexity of the environmental space, limit the number of dimensions involved, and reduce collinearity.

Model averaging: The medians of distinct replicates of Maxent models created with different parameter settings and sets of variables were used to summarize all results.

#### Workflow

Model workflow: Model calibration was done within a 500 km buffer around species' occurrences. To consider uncertainty deriving from specific treatments of occurrence records and environmental predictors, we calibrated models via four distinct schemes: (1) raw variables and distance-based thinned occurrences, (2) PCs and distance-based thinned occurrences, (3) raw environmental variables and country-density thinned occurrences, and (4) PCs and country-density thinned occurrences. For each scheme, we calibrated models five times, each time randomly selecting 50% of occurrences for training models, and using the remaining records for testing. We assessed model performance using partial ROC (for statistical significance), omission rates (E = 5%, for predictive ability), and Akaike Information Criterion corrected for small sample sizes (AICc). We selected models with delta AICc ≤2 from among those that were statistically significant and had omission rates below 5%. We created models with the selected parameter values, using all occurrences after the corresponding thinning process, with 10 bootstrap replicates, cloglog output, and model transfers to the world using three types of extrapolation (free extrapolation, extrapolation and clamping, and no extrapolation). As a final evaluation step, we tested whether each replicate of the selected models was able to anticipate the known invasive records of the species in the Americas (British Columbia, Canada; Washington, USA). We created two types of consensus: (1) a median of the medians obtained for each parameterization, and, (2) the sum of all suitable areas derived from binarizing each replicate using a modified least presence (5% omission) threshold. We used the mobility-oriented parity metric (MOP) to detect areas where strict or combinational extrapolation risks could be expected, and we used those areas to trim our binary results. Areas detected as suitable (excluding areas of strict extrapolation) were used as the base for simulation of potential invasion patterns of this hornet in North America using a cellular automaton dynamic model. Simulations were done for results of all four schemes of data processing.

##### Software

Software: Maxent 3.4.1; R 3.6.2 (packages: bam, biosurvey, ellipsenm, kuenm, ntox, raster, rgdal, and rgeos)

Code availability: https://github.com/townpeterson/vespa

Data availability: http://hdl.handle.net/1808/30602

#### Data

##### Biodiversity data

Taxon names: *Vespa mandarinia*

Taxonomic reference system: N/A

Ecological level: species

Data sources: Occurrence data for *V. mandarinia* were downloaded from the Global Biodiversity Information Facility database (GBIF; https://www.gbif.org/).

Sampling design: Spatial thinning and maximum density per country thinning

Sample size: 18 (country-density thinned occurrences); 49 (distance-based thinned occurrences)

Clipping: east Asia (native range of distribution of this species)

Scaling: N/A

Cleaning: We kept records from the species' native range separate from non-native occurrences facilitated by human introduction. We cleaned occurrences from the native distribution following Cobos et al. (2018) by removing duplicates and records with doubtful or missing coordinates. To avoid model overfitting derived from spatial autocorrelation and overdominance of specific regions due to sampling bias, we thinned these records spatially in two ways: by geographic distance and by density of records per country. In the first case (distance-based thinning), we excluded occurrences that were <50 km away from another locality. In the second thinning approach (country-density thinning), we randomly reduced numbers of occurrences in countries with the densest sampling, namely Japan, Taiwan, and South Korea (from 30, 6, and 5, to 6, 2, and 2 occurrences, respectively), to match an approximate reference density of India, Nepal, and China.

Absence data: N/A

Background data: A buffer of 500 km around species records for spatial delimitation of calibration areas. Records from 1990 onwards.

Errors and biases: N/A

#### Predictor variables

Predictor variables: We used two types of predictors: 1) Six raw bioclimatic variables selected based on correlation levels and species natural history criteria: isothermality (BIO3), maximum temperature of warmest month (BIO5), minimum temperature of coldest month (BIO6), temperature annual range (BIO7), specific humidity of most humid month (BIO13), and specific humidity of least humid month (BIO14). 2) The first four principal components (PC) axes of a PCA analysis done with 15 variables (all variables of MERRAclim but Bio8, Bio9, Bio18, and Bio19), as they explained 97.9% of the cumulative variance.

Data sources: MERRAclim database (Vega, Pertierra & Olalla-Tárraga, 2018)

Spatial extent: 72.856955, 148.618463, 18.190443, 48.987058 (xmin, xmax, ymin, ymax)

Spatial resolution: 10’ resolution (~18 km at the Equator)

Coordinate reference system: +proj=longlat +datum=WGS84 +no_defs

Temporal extent: 2000-2010

Temporal resolution: N/A

Data processing: N/A

Errors and biases: N/A

Dimension reduction: N/A

#### Transfer data

Spatial extent: -180, 180, -90, 83.623596 (xmin, xmax, ymin, ymax)

Spatial resolution: 10’ resolution (~18 km at the Equator)

Temporal extent: 2000-2010

Temporal resolution: N/A

#### Model

##### Variable pre-selection

Variable pre-selection: We excluded four of the "bioclimatic" variables because they are known to contain spatial artifacts as a result of combining temperature and humidity information (Escobar et al., 2014): mean temperature of most humid quarter, mean temperature of least humid quarter, specific humidity mean of warmest quarter, and specific humidity mean of coldest quarter. The 15 variables remaining were masked to an area for model calibration. These 15 variables were submitted to a principal component analysis (PCA) to reduce dimensionality and multicollinearity. To select a set of raw variables, we reduced them to a subset with Pearson's correlation coefficients ($r$) ≤ 0.85, choosing the most biologically relevant or interpretable variables based on our knowledge of AGH natural history. The PCA was calibrated using environmental variation across the M area, and transferred to the whole world.

##### Multicollinearity

Multicollinearity: We did a principal components analysis to reduce multicollinearity. We also used Pearson's correlation coefficient values to select a subset of non-correlated variables.

##### Model settings

Maxent: Feature Set (eight feature classes (lq, lp, lqp, qp, q, lqpt, lqpth, lqph, where l is linear, q is quadratic, p is product, t is threshold, and h is hinge)), Regularization Multiplier Set (10 regularization multiplier values (0.10, 0.25, 0.50, 0.75, 1, 2, 3, 4, 5, 6)), Preictor Set (All combinations of more than 2 predictors of the first 4 PCs. All combinations of more than 2 variables of the six selected biolimatic predictors.)

##### Model estimates

Coefficients: Median

Parameter uncertainty: N/A

Variable importance: N/A

##### Model selection - model averaging - ensembles

Model selection: N/A

Model averaging: N/A

Model ensembles: N/A

##### Analysis and Correction of non-independence

Spatial autocorrelation: N/A

Temporal autocorrelation: N/A

Nested data: N/A

#### Assessment

##### Performance statistics

Performance on training data: True positive rate, AIC, AUC

##### Plausibility check

Response shapes: No plausibility checks conducted
